# The dynamics of ciliogenesis in prepubertal mouse meiosis reveal new clues about testicular maturation during puberty

**DOI:** 10.1101/2025.06.19.660582

**Authors:** I Pérez-Moreno, P López-Jiménez, H Zapata, S Pérez-Martín, M López-Panadés, B Barbeito, J Urtasun-Elizari, I Roig, FR Garcia-Gonzalo, J Page, R Gómez

**Affiliations:** Unidad de Biología Celular, Departamento de Biología, Universidad Autónoma de Madrid, 28049 Madrid, Spain; MRC Laboratory of Medical Sciences London W12 0NN UK; Centro de Investigación Biomédica en Red en Enfermedades Cardiovasculares (CIBERCV), 28029 Madrid, Spain; Department of Pharmacology, School of Medicine, Instituto de Investigación Sanitaria Gregorio Marañón, Universidad Complutense, 28040 Madrid, Spain; Genome Integrity and Instability Group, Institut de Biotecnologia i Biomedicina, Universitat Autònoma de Barcelona, 08193 Cerdanyola del Vallès, Spain; Cytology and Histology Unit, Department of Cell Biology, Physiology and Immunology, Universitat Autònoma de Barcelona, 08193 Cerdanyola del Vallès, Spain; Departamento de Bioquímica, Facultad de Medicina, Universidad Autónoma de Madrid, 28029 Madrid, Spain; Departamento de Enfermedades Raras, Instituto de Investigaciones Biomédicas Sols-Morreale (IIBM), CSIC-UAM, Madrid, Spain; CIBER de Enfermedades Raras (CIBERER), Instituto de Salud Carlos III, Madrid, Spain

**Keywords:** meiosis, cilia, centrosome, cysts, prepuberal mice, testis

## Abstract

The primary cilium, a solitary and non-motile extension of the plasma membrane, has recently been identified in adult male mouse spermatocytes. However, very little is known about when these cilia emerge during testicular maturation and what their function is. In the context of fertility establishment during puberty, this study investigates the dynamics of ciliogenesis in prepubertal mouse spermatocytes. Our findings reveal that primary cilia are not an intrinsic feature of spermatocytes during the first wave of meiosis, which initiates at 8 days post-partum (dpp). Instead, cilia begin polymerizing at 20 dpp, after first meiotic wave has been completed, and are present in spermatocytes across all stages of prophase I. Thus, no direct correlation between cilia polymerization and the initiation of synapsis or desynapsis was found, although chemical ablation of cilia may delay DNA repair during prophase I. Typical adult cilia features, which are shorter and restricted to zygotene spermatocytes, are settled upon acquisition of sexual maturity. This study also highlights that the emergence of ciliated spermatocytes in prepuberal mice coincides with the onset of flagellogenesis, hinting at a potential link between the regulation of the formation of both types of axonemes within the tissue developing environment. Proteomic analyses further identify temporal regulators of axoneme assembly, providing valuable targets for future research to unravel the molecular pathways underlying ciliogenesis, flagellogenesis, and their roles in spermatogenesis. We explored distinct regulatory mechanisms of ciliogenesis during the first meiotic wave and found that Aurora kinase A (AURKA) is a critical regulator of cilia disassembly during late diplotene, with evidence suggesting that centrosome migration and cilia depolymerization are mutually exclusive events during meiosis. In summary, this study provides the first detailed characterization of primary cilia dynamics during early testicular maturation in mice, revealing their spatiotemporal regulation, candidate molecular mediators, and potential roles during meiosis. These findings lay the groundwork for understanding the physiological relevance of meiotic cilia in spermatogenesis and testicular development.

## INTRODUCTION

Cilia and flagella are microtubule (MT)-based plasma membrane projections that function as motile or sensory structures. Motile cilia and flagella have a well-established structure, typically exhibiting an axoneme with a 9+2 arrangement, with nine peripheral MT doublets connected to each other by dynein arms, and a central MT pair bound to peripheral doublets by radial spokes. This organization, along with the association of motor proteins, enables motile cilia and flagella to generate rhythmic movements that result in cell locomotion, extracellular fluid displacement or spermatozoan propulsion ^1-3^. In contrast, primary cilia serve as signalling organelles, detecting mechanical, optical, or chemical stimuli ^4^. These primary cilia exhibit unique features. First, they are solitary, with only one present per cell. Second, they are transient structures, only present during the G1/G0 phases of the somatic cell cycle. Third, they possess a 9+0 axoneme configuration. And finally, they lack motility since their axoneme does not contain motor proteins ^5^.

Despite functional and structural differences, motile cilia, primary cilia, and flagella share numerous molecular and architectural components. In all cases, the axoneme arises from the mother centriole of a membrane-docked centrosome, also referred to as the basal body. The mother centriole presents distal and subdistal appendages that facilitate the docking to the membrane ^6^. Additionally, they share several components, such as MTs with acetylated tubulin (AcTub) or the ADP Ribosylation Factor Like GTPase 13B (ARL13B) ^7-9^. Notably, the assembly, maintenance, and functions of both motile and nonmotile cilia strictly depend on intraflagellar transport (IFT) proteins ^10,11^. As cilia lack the translational machinery for protein synthesis, IFT serves as the sole mechanism for trafficking the essential components into the ciliary compartment required for cilia assembly, growth and functions. Interestingly, several IFT proteins, including IFT88, are also localized to centrosomes at mitotic spindle poles ^12,13^. Beyond their role in ciliated cells, IFT88 and related proteins, partner with Dynein1-driven complexes to transport peripheral MT clusters and MT-nucleating proteins to spindle poles, facilitating astral MT formation and proper spindle orientation ^14^.

Over the past century, research has uncovered the diverse functions of primary cilia in mammalian tissues, including roles in mechanosensing, vision, and chemical signal transduction. Playing these functions, cilia can detect a wide range of stimuli—including blood flow, light, hormones, and growth factors—underscoring their importance in cellular signalling ^4,15-17^. Given these roles, disruptions in ciliary structure or signalling pathways due to mutations in cilia-related genes result in a spectrum of diseases known as ciliopathies ^18^. These disorders can affect ciliary assembly and maintenance or, in the case of primary cilia, disrupt intracellular signalling pathways ^17,19,20^.

Advancements in our understanding of primary cilia have uncovered that ciliary assembly (ciliogenesis) and disassembly are dynamically regulated in coordination with the cell cycle ^21^. In somatic cells, primary cilia form in G1/G0 and are reabsorbed in G2, thereby freeing centrioles and enabling them to function as Microtubule Organizing Centers (MTOCs) at the spindle poles during cell division ^22-24^. Key regulators of primary cilium disassembly appear to include many well-known master controllers of mitotic progression ^25,26^. Among these, the mitotic Aurora A kinase (AURKA) ^27^ has emerged as a crucial regulatory factor for somatic primary cilium disassembly. Activation of AURKA in the context of primary cilium resorption is mediated by a mechanism involving the phosphorylation and activation of deacetylase HDAC6 ^28,29^.

Gametogenesis is a fundamental biological process responsible for generating gametes (oocytes and spermatozoa). Spermatogenesis, in particular, comprises at least three functional processes: i) proliferation of spermatogonia (stem germ cells), ii) meiosis, a specialized cell division that generates haploid cells (spermatids) from the diploid spermatogonia, and iii) spermiogenesis, i.e., the morphological maturation of spermatids into mature spermatozoa, which includes the formation of the flagellum ^30^. In mammals, spermatogenesis occurs in the seminiferous epithelium of the testis, a highly specialized environment that allows nonsynchronous and continuous spermatogenesis. A key participant of spermatogenesis are the Sertoli cells, which are the supporting cells of the seminiferous epithelium and play a crucial role in providing the structural and hormonal environment for spermatogenesis ^31^. Another key feature of the seminiferous epithelium is the fact that spermatocytes and spermatids are interconnected via cytoplasmic bridges, forming large cysts ^32^, which facilitates the biochemical and physiological coordination of cells within the epithelium ^33,34^.

The testis undergoes profound physiological and morphological transformations from fetal development to adulthood, driven by complex hormonal and molecular adjustments that promote the maturation of spermatogonia and the initiation of spermatogenesis ^35-37^. Remarkably, the testis is one of the few organs that establishes most of its cell types, functions, and physiology after birth ^38^. Puberty marks a critical period during which the inaugural wave of complete spermatogenesis initiates with spermatogonial differentiation and culminates in the production of the first spermatozoa. This initial wave not only establishes male fertility but also sets the foundation for continuous sperm production throughout the reproductive lifespan. Hence, as the stage when fertility is established, puberty provides a critical window to investigate key events underlying spermatogenesis.

In recent years, primary cilia have been observed in several somatic cell types of the testis and epididymis, including Sertoli, peritubular myoid or epididymal cells ^39^. Moreover, the presence of cilia in germ cells has now been confirmed in various species, including *Drosophila melanogaster* spermatocytes ^40^, zebrafish oocytes and spermatocytes ^41,42^, and mouse spermatocytes ^43^. These findings challenge the long-standing paradigm that primary cilia must be disassembled prior to cell division. Indeed, the presence of the cilium seems to be critical for chromosome dynamics at the beginning of meiosis in zebrafish, and its disorganization can compromise meiosis outcome ^41^.

These findings open intriguing questions on the relationship between gametogenesis and ciliogenesis that call for a deeper characterization. Nevertheless, reports studying the extent and features of primary cilia in mammalian spermatocytes are scarce. In a previous study we found that in adult mice cilia are prominent and appear predominantly in zygotene-stage spermatocytes, suggesting that cilia can play an important role in the physiological and functional coordination of spermatocytes within the seminiferous tubules ^43^. In contrast, it has been suggested that in prepubertal mouse spermatocytes cilia may be much shorter ^41^. This makes us suspect that there may be differences in the development and/or regulation of cilia formation in the first spermatogenesis wave compared to adults. To address this question, we have performed a systematic study of primary cilia morphogenesis in prepubertal mice, from day 8 until day 60 post-partum. Our results discovered that the timing of cilium formation in spermatocytes is temporally linked to flagellum development in spermatids. These findings suggest an unanticipated coordination in the formation of axonemal structures during the establishment of fertility in puberty, pointing to a broader developmental program governing ciliogenesis in the prepuberal male germline, which may depart from the formation of cilia in adult spermatogenesis.

## RESULTS

### The first wave of spermatogenesis is completed 28 days *post partum*

Given the discrepancies previously reported about ciliogenesis in mouse spermatocyte ^41,43^, we hypothesized about the possibility that cilia morphogenesis in spermatocytes is a progressive process occurring during puberty. To investigate this process, we first characterized the progression of the first spermatogenesis wave in mice, as striking discrepancies in timeline were found across earlier studies ^44-48^. To this end, testes from prepubertal, lactating C57BL/6 male mice were collected, measured, and weighed daily between 8 and 30 dpp. In addition, we analysed non-lactating animals between 30 dpp and 60 dpp at weekly intervals. As expected, testis weight increased steadily with age, reaching adult-like weight between 45 and 60 dpp (Fig. 1-I). Stage 8 dpp marks the appearance of the first meiotic spermatocytes derived from self-renewing undifferentiated spermatogonia ^49^. At this age, the seminiferous tubules exhibited a small diameter and contained only spermatogonia and primary spermatocytes (Fig. 1-II A). The diameter of the seminiferous tubules increased progressively, and round (immature) spermatids were first observed in testis sections at 21 dpp (Fig. 1-I B-D). By 30 dpp, elongated spermatids and mature spermatids were observed in the lumen of the seminiferous tubules; however, these animals were still considered prepubertal, evidenced by the absence of mature spermatids or spermatozoa in epididymal sections (Fig. 1-I E and E′). Fully mature spermatozoa were first detected in the epididymis at 38 dpp (Fig. 1-I F and F′). From 45 dpp onwards, the testis was fully developed, with seminiferous tubules growing towards typical adult diameter and epididymal sections progressively filling with spermatozoa (Fig. 1-I G and G′). At 60 dpp, epididymis is densely packed with spermatozoa (Fig. 1-I H and H′).

**Figure 1.**
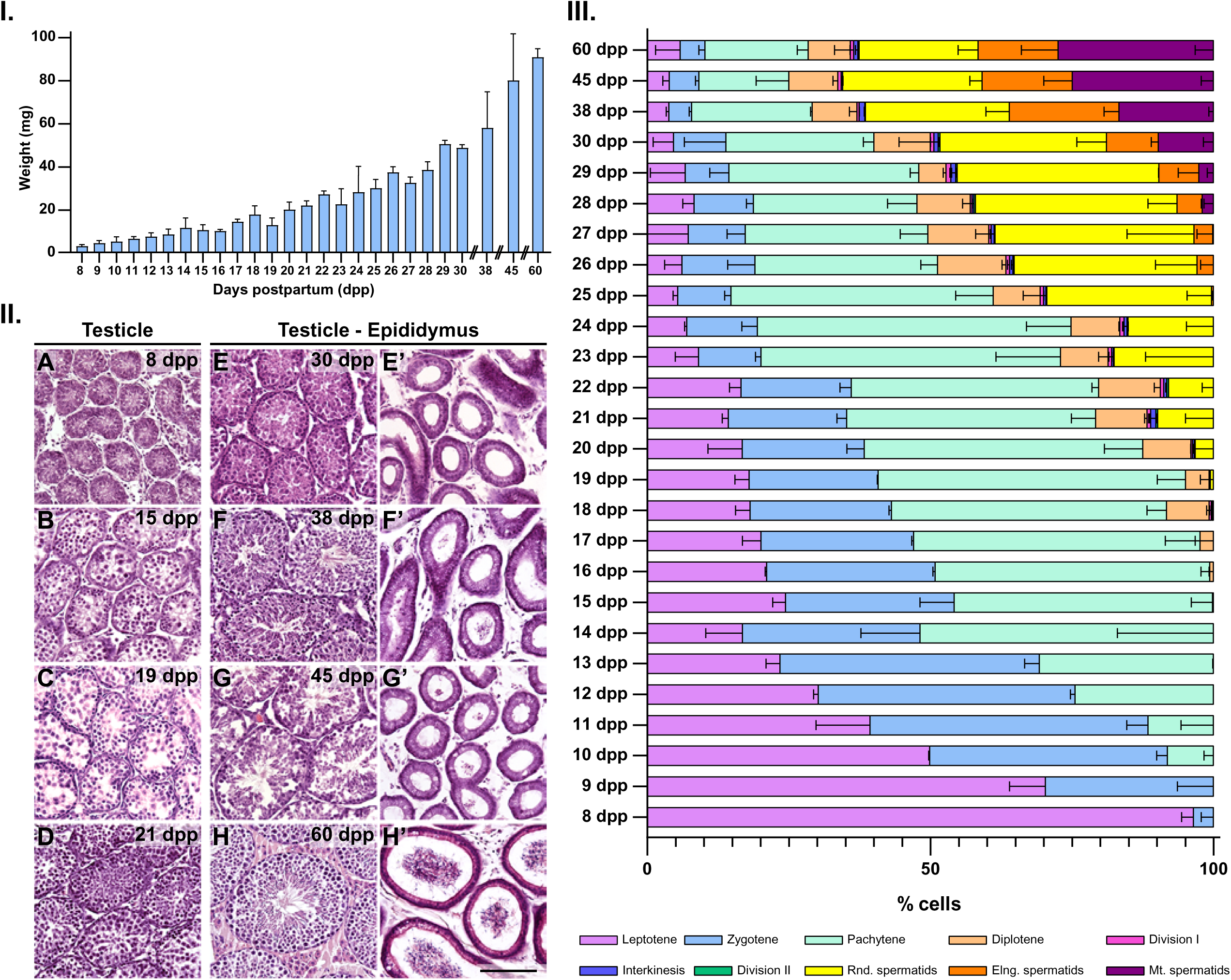
The first spermatogenesis wave in prepuberal mice. **I.** Graphical representation of the weight of the testis in animals in the age range 8 to 60 days post partum (dpp) (two biological replicates from 8 to 22 dpp and three biological replicates from 23 to 60 dpp) **II.** Cross sections of the H&E-stained testis or testis and epididymis from 8, 15, 19, 21, 30, 38, 45 and 60 dpp mice. Scale bar represents 100 µm. **III.** Distribution of the meiosis stages in animals in the age range 8 to 60 dpp. The analysis quantifies stages at prophase I (leptotene, zygotene, pachytene and diplotene), division I (metaphase I, anaphase I and telophase I), interkinesis, division II (metaphase II, anaphase II and telophase II), and spermatids (round/immature, elongated, and mature) (two biological replicates for 8 and 9 dpp n=250 cells, and two biological replicates for 10 to 60 dpp n=2000 cells).

We later characterized the proportion of spermatocytes at different stages during testis maturation (Fig. 1-III), analysing the presence and localization of SYCP3 and SYCP1 proteins of the synaptonemal complex, along with γH2AX (a marker for DNA double-strand breaks -DSBs-) in squashed seminiferous tubules of mice between 8 and 60 dpp (Supplementary Fig. 1). We recorded the following stages: prophase I (leptotene, zygotene, pachytene, diplotene, diakinesis), first meiotic division (metaphase I, anaphase I and telophase I), interkinesis, second meiotic division (metaphase II, anaphase II and telophase II). We also recorded spermatids at different stages of spermiogenesis (round spermatids, elongated spermatids and mature spermatids). The most relevant findings indicate that at 8 dpp, 96.49% of spermatocytes are at the leptotene stage. Pachytene spermatocytes first appeared at 10 dpp, while the first and second meiotic divisions—as well as the interkinesis stage in between them—were observed for the first time at 18 dpp (Fig. 1-III). Round spermatids were detectable from 19 dpp, with mature and flagellated late spermatids first appearing at 28 dpp. The proportion of mature spermatids increased progressively between 28 and 45 dpp. Based on our data, and in agreement with the histological analysis (Fig. 1-II), 45 dpp can be considered the onset of adult testicular function, as the ratio of spermatids stabilizes from this point through 60 dpp and beyond. (All values for Figure 1 are included in Supplementary Table 1).

### Ciliogenesis simultaneously emerges across all stages of prophase I spermatocytes during the initial spermatogenesis wave

We then aimed to identify the presence of cilia in testicular cells. For this purpose, we double immunolocalized SYCP3 and Acetylated Tubulin (AcTub), a marker of somatic ^50^ and meiotic ^43^ primary cilium. Our results show that mice between 8 to 19 dpp did not present primary cilium in any of the stages of prophase I. AcTub labeling was only detected at the centrosomes, as two small and rounded signals close to each other and to the cell nucleus, corresponding to the pair of centrioles (Fig 2-I A-D). In contrast, prepuberal individuals from 20 dpp onwards presented a small but significant proportion of spermatocytes at prophase I showing a ciliary axoneme labelled with AcTub (Fig 2-I Á-D′). Ciliated spermatocytes at 20 dpp represented 2.34% of all prophase I cells, increasing to approximately 5% in mice at 21 and 22 dpp, and then progressively decreasing until 30-45 dpp, when the percentage of prophase I ciliated spermatocytes (0.43%) is similar to adulthood (Fig. 2-II A), being consistent with previously reported data for adult meiosis^43^. Unexpectedly, cilia were observed in spermatocytes at different stages of prophase I. This is in stark contrast to adult spermatogenesis, in which a primary cilium is only present in zygotene spermatocytes ^43^. Instead, in prepuberal mice, most ciliated spermatocytes were at pachytene stage, followed by diplotene and zygotene, while leptotene stage showed lower proportions of ciliated cells (Fig. 2-II B). The sudden appearance of the primary cilium in different prophase I stages at 20 dpp, suggests that this is a simultaneous process in all prophase I spermatocytes at this developmental phase. Spermatocytes during later stages of the first and second meiotic division, or at interkinesis, did not present cilia.

**Figure 2.**
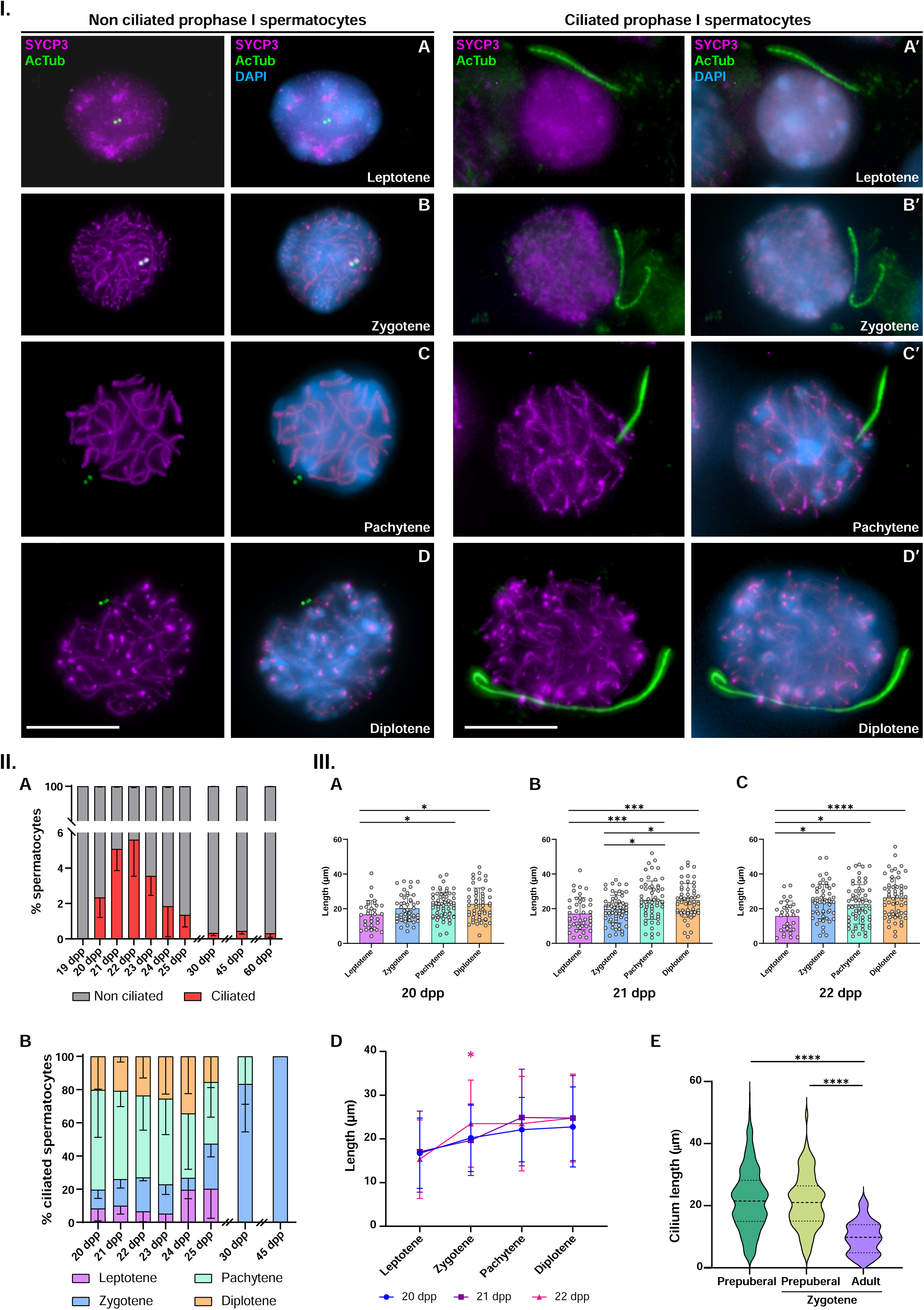
Cilia emerge across all stages of prophase I in spermatocytes during the first spermatogenesis wave. **I. Cilia appear in spermatocytes at leptotene, zygotene, pachytene and diplotene. A-D.** Double immunolabelling of acetylated Tubulin (AcTub) (green) and synaptonemal complex protein 3 (SYCP3) (magenta) with chromatin stained with DAPI (blue) on squashed WT mouse spermatocytes at (A and Á) Leptotene, (B and B′) Zygotene, (C and Ć) Pachytene and (D and D′) Diplotene. All images are z-projection of 30-60 frames of stack images of each cell. Scale bars represent 10 µm. **II. Cilia quantification. A.** Graphical representation of the quantification of ciliated spermatocytes at ages between 20 and 60 days post partum (dpp). **B.** Graphical representation of the ciliated spermatocytes at different stages of prophase I for the age frame 20-60 dpp (n=1500, three biological replicates except for 45 dpp in which n=1100, two biological replicates). **III. Cilia length quantification. A-C.** Graphical representation of cilia length at different stages of prophase I in (A) 20dpp, (B) 21 dpp and (C) 22 dpp prepuberal animals. Data represents mean ± SD, **** p < 0.0001, One-way ANOVA, a minimum of 30 spermatocytes were quantified per stage (three biological replicates). **D.** Graphical representation of cilia length at different stages of prophase I in a combined analysis for 20-22 dpp mice. **E.** Graphical representation of the global cilia length in all stages of prophase I spermatocytes, and specifically in zygotene, in prepuberal versus zygotene spermatocytes in adult mice. Data represents mean ± SD, **** p < 0.0001, Kruskal-Wallis test (n=360, at least 30 cilia per prepuberal spermatocyte for each stage, and n=65, at least 20 zygotene adult spermatocytes. Both prepuberal and adult analysis has been made in three biological replicates).

Next, we measured the length of the primary cilium for each meiotic stage and age group. The length of the cilium was found to vary depending on the meiotic stage, reaching its maximum length at pachytene and diplotene stages. No differences were found in the length of the primary cilium between different ages, except between 21 and 22 dpp in zygotene cells (Fig. 2-III A-D). Notably, prepuberal meiotic cilia are significantly longer (21,96 μm, rank: 3 to 55 μm) than adult cilia (10 μm) ^43^, even when comparing only ciliated spermatocytes at zygotene (Fig. 2-III E).

### Cilia structure in mouse spermatocytes

Overall, the results presented above indicate that the length and the cell stage at which primary cilia are shaped clearly differ in prepuberal compared to adult mice. To ascertain whether other features are shared or not, we first analyzed the location and dynamics of additional markers for cilia and centrosome.

Cilia in prepuberal spermatocytes showed ARL13b and AcTub co-labelling (Fig 3-I A), as previously reported for adult ciliated spermatocytes (López-Jiménez et al., 2022). We also found that meiotic prepuberal cilia arise from the maternal centrioles, labelled with CEP164 in all stages of prophase I (Fig 3-I B-D). Moreover, we detected that cilium in prepuberal spermatocytes contain IFT component IFT88, which faintly labels the cytoplasm and the ciliary axoneme at all stages of prophase I (Fig 3-I E-G and E′-G′). This is the first time that this protein has been shown in spermatocytes and suggests active IFT in these cilia. Interestingly, IFT88 was also present at centrosomes in ciliated (Fig. 3-I E′-G′ and Fig. 3-II C) and non-ciliated spermatocytes (Fig 3-II A-E). When co-detected with CETN3, IFT88 labeled the maternal centriole in non-ciliated spermatocytes at early prophase I (Fig 3-II A-B) and started to appear in both duplicated centrosomes once their migration starts at late diplotene (Fig 3-II D). When reaching metaphase I, IFT88 is present at both bipolar centrosomes of the dividing spermatocyte (Fig 3-II E). Thus, the organization and composition of primary cilia in prepuberal spermatocytes seems similar to the ones described in their adult counterparts, and also in cilia of somatic cells.

**Figure 3.**
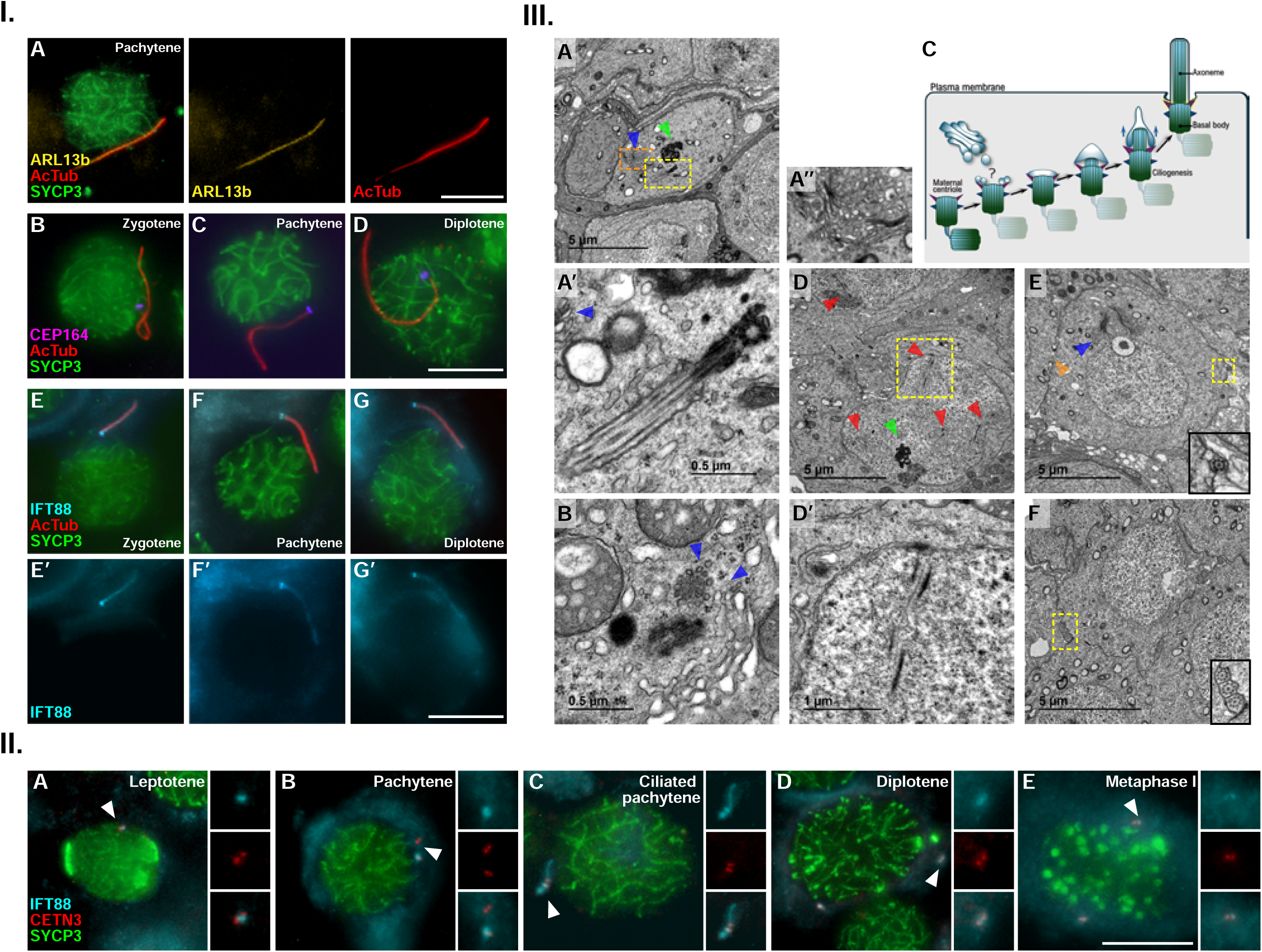
Ultrastructure and markers of prepuberal meiotic cilia. **I. Distribution of ARL13B, CEP164 and IFT88 in ciliated spermatocytes. A-G.** Triple immunolabelling of acetylated Tubulin (AcTub) (red), synaptonemal complex protein 3 (SYCP3) (green) and (A) ARL13b (yellow), (B-D) CEP164 (magenta) or (E-G) IFT88 (cyan) in mouse spermatocytes during prophase I (zygo-tene, pachytene and diplotene). All images are z-projection of 30-60 frames of stack images of each cell. Scale bar in G′ represents 10 µm. **II. Distribution of IFT88 at meiotic centrosomes. A-E.** Triple immunolabelling of IFT88 (cyan), Centrin 3 (CETN3) (red) and synaptonemal complex protein 3 (SYCP3)(green) in mouse spermatocytes during prophase I (leptotene, pachytene and diplotene) (A-D) and Metaphase I (E). White arrowheads point to selected centrosome for the amplified image. All images are z-projection of 30-60 frames of stack images of each cell. Scale bar represents 10 µm. **III. Ultrastructure of axonemes in prepuberal meiosis. A.** Electron microscopy images of a ciliated spermatocyte. Amplified images of the cilia (Á) and the Golgi apparatus (Á′). **B.** Centrosomes with Golgi derived vesicles in the maternal centriole. **C.** Schematic representation of intracellular ciliogenesis pathway. **D.** Prophase I spermatocyte. **E-F.** Mid spermatids presenting preacrosomic vesicle. Enhanced images of the flagella are offered. Blue arrowheads point to Golgi apparatus vesicles. Red arrowheads point to fragments of synaptonemal complex. Green arrowheads point to nucleolus. Orange arrowheads point to intercellular bridge between spermatids. Scale bars are included in each image.

To gain a more complete analysis of the cilia organization, we further analyzed their ultrastructure in prepuberal spermatocytes under transmission electron microscopy (TEM) (Fig 3-III). The meiotic cilia were observed in spermatocytes that present typical features of prophase-I, like a well-developed nucleolus and fragments of synaptonemal complex (Fig 3-III A and D-D′ and Supplementary Fig 2). Cilia axoneme emerges from one of the centrioles and typically lack a central pair of MT. In contrast, flagella observed in early spermatids, already present in 21 dpp mice, are distinguishable for having a 9+2 axonemal organization and a clear preacrosomal vesicle near the nucleus (Fig 3-III E). We observed that meiotic cilia did not polymerize attached to the plasma membrane. Instead, they might emerge from the mother centriole of the centrosome, while it is still located internally in the cell, close to the nuclear envelope (Fig 3-III. A and A′). Our analysis suggests that cilia formation may follow an intracellular pathway, in which vesicles originating from the Golgi apparatus (Fig 3-III. A′′) are transported to the region above the maternal centriole, where they subsequently fuse (Fig 3-III B). Then, MT axonemes begin to polymerize within the cytoplasm and subsequently extend beyond the cell membrane into the extracellular space (see schematic diagram in Fig 3-III C). This pathway has been previously described for several somatic tissues ^51-54^. Taking advantage of the resolution of TEM, we have additionally analysed selected meiotic centrosomes at different stages of prophase I: at leptotene, prior to duplication, and at zygotene, after duplication (Supplementary Fig 3 A-C). Centrosomes in spermatocytes at these stages have been rarely documented in recent decades, following the classic studies ^55^.

Our electron microscopy analysis also showed that prophase I spermatocytes in prepuberal mice are interconnected via cytoplasmic bridges and share cytoplasmic components, including mitochondria (Fig. 4-I A). These bridges, partially composed of the Testis-expressed protein 14 (TEX14), are present from the onset of the first meiotic wave and connect spermatocytes within the same cyst (Fig 4-I B), a feature also observed in adult mice ^43^. The continuity of the shared cytoplasm in cyst of spermatocytes is demonstrated in electron microscopy images (Fig 4-I A), and with the immunostaining of the germ specific cytoplasmic marker VASA ^43,56^ (Fig 4-I B). This histological organization is conserved in humans, where ciliated spermatocytes are similarly observed at the base of seminiferous tubules (Fig 4-II and Supplementary Fig 4). These findings indicate that cysts of interconnected spermatocytes are already established during the first meiotic wave persisting into adulthood in mouse, and at least in adulthood in humans. The evolutionary conservation of this histological organization highlights its plausible fundamental role in mammalian spermatogenesis.

**Figure 4.**
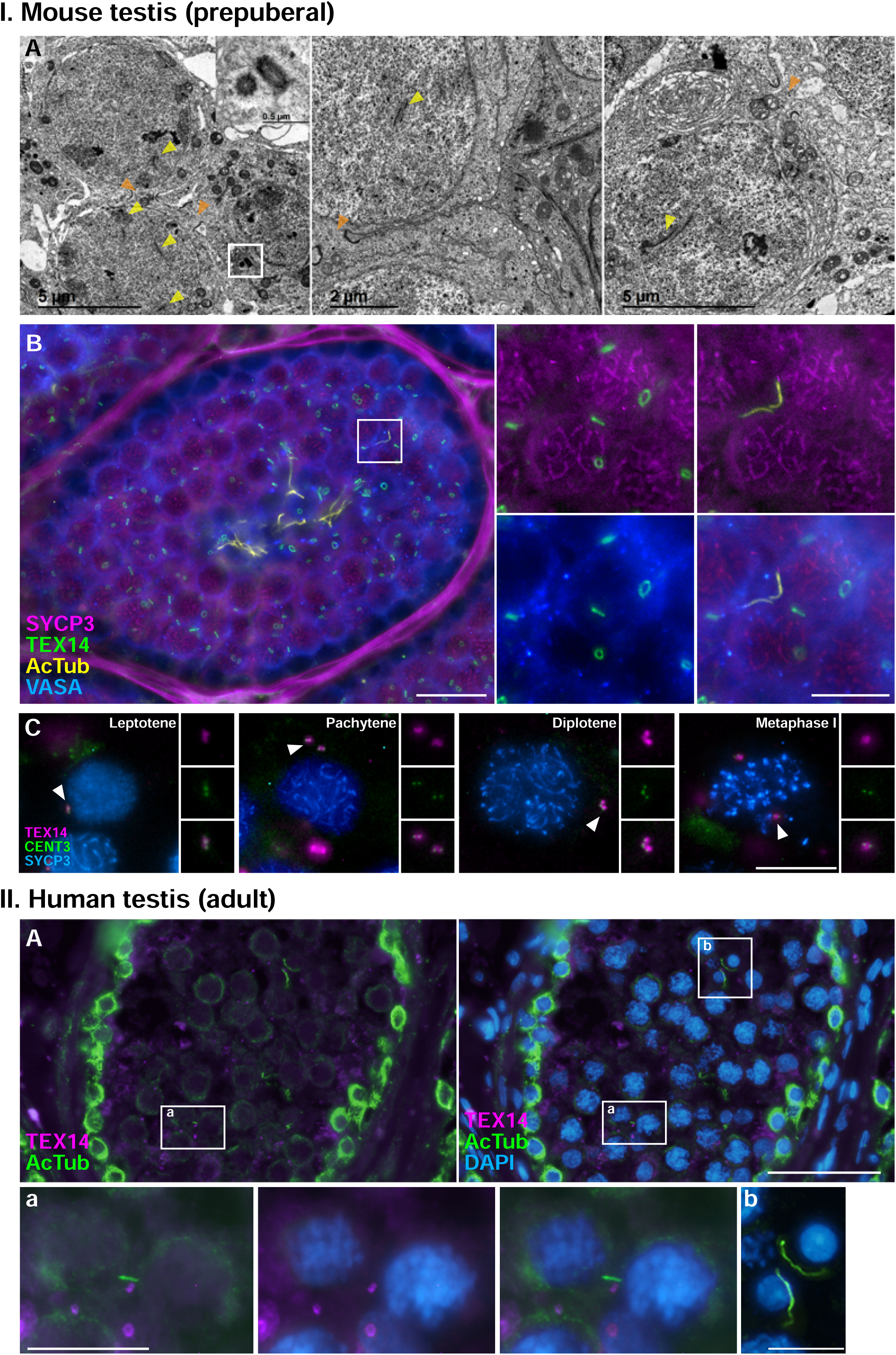
Testicular tissue presents spermatocyte cysts in prepuberal mice and adult humans. **I. Mouse spermatocytes are connected in cysts from the first round of spermatogenesis. A.** Electron microscopy images of spermatocytes at prophase I presenting fragments of synaptonemal complex (yellow arrowheads) which are connected by intercellular bridges (orange arrowheads). The centrosome of one of the spermatocytes is enhanced in an amplified image. Scale bars are included in each image. **B.** Quadruple immunolabelling of synaptonemal complex protein 3 (SYCP3) (magenta), testis-expressed protein 14 (TEX14) (green), acetylated Tubulin (yellow) and the germ specific cytoplasmic marker VASA 41,54 (blue) in testis cryosection of a 21 dpp mouse. Scale bars represent 20 µm for section images and 10 µm for amplified sections. **C.** Triple immunolabelling of (TEX14) (magenta), Centrin 3 (CETN3) (green) and SYCP3 (blue) in squashed spermatocytes at prophase I (leptotene, pachytene and diplotene) and metaphase I. Scale bar represents 10 µm. **II. Human spermatocytes are connected in cysts and present cilia. A.** Double immunolabelling of TEX14 (magenta) and acetylated Tubulin (green) in adult human testis cryosection. Chromatin is stained with DAPI (blue). Selected enhanced images correspond to a ciliated spermatocyte and a flagellated early spermatid. Scale bars represent 50 µm in A and 10 µm in (a) and (b).

### Cillia and flagella appearance are temporarily linked during prepuberal meiosis

As corroborated by both light and electron microscopy, the first appearance of cilia in spermatocytes occurs concurrently with the formation of the first spermatids. Thus, we asked whether these two events were occurring within the same seminiferous tubules. To this end, it was important to maintain the histological organization of cells, so we performed this analysis in mouse testis sections at 19, 20, 21 and 30 dpp, studying the distribution of AcTub in combination with SYCP3 (Fig. 5). In agreement with our results in squashed spermatocytes, ciliated spermatocytes are not observed in 19 dpp mice (Fig. 5 A), but were detected at 20 dpp (Fig. 5 B). Interestingly, these same tubules already included immature round spermatids, that have initiated the formation of the flagellum, which projects towards the lumen of the seminiferous tubule (Fig. 5 B -B’’). This histological organization is also observed in prepubertal mice at 21 dpp, when elongation of flagella had begun (Fig. 5 C-C’’), and at 30 dpp, were the percentage of ciliated prophase I spermatocytes is similar to adulthood (Fig. 2-II A). By this stage, a lumen filled with mature spermatids resembling adult testis histology was observed (red arrowhead in Fig. 5 D-D’’), as shown in Figure 1.

**Figure 5.**
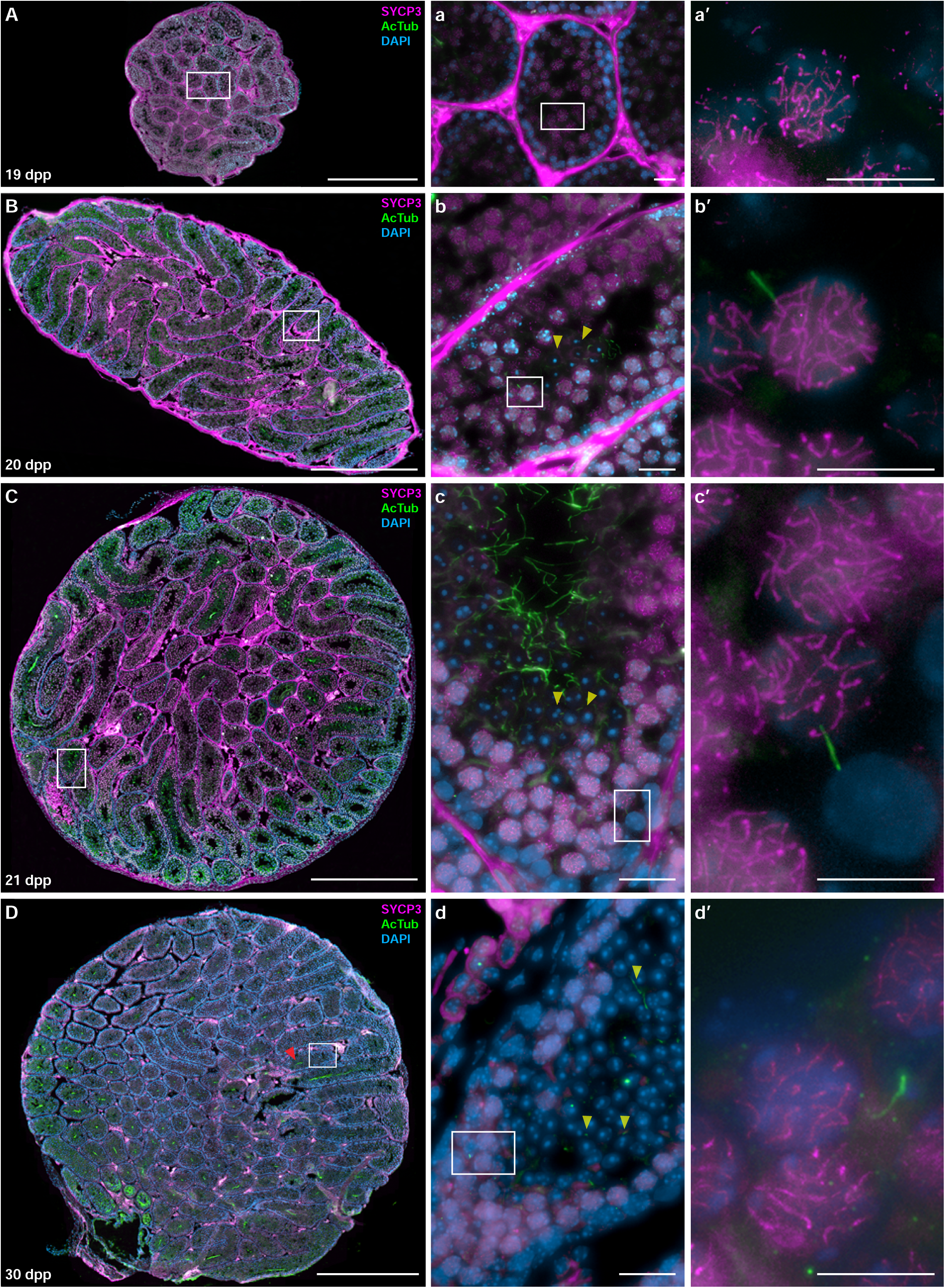
Cillia and flagella dynamics are correlated during prepuberal meiosis. **A-D.** Double immunolabelling of acetylated Tubulin (green) and synaptonemal complex protein 3 (SYCP3) (magenta) in whole testis cryosection of 10 dpp (A), 20 dpp (B), 21 dpp (C) and 30 dpp (D) mice. Chromatin is stained with DAPI (blue). Enhanced images correspond to selected seminiferous tubules (a-d), where ciliated spermatocytes are identified in the bases of the tubule after 20 dpp (b′-d′). Early spermatids are indicated with yellow arrowheads, and late spermatids are indicated with red arrowheads. Scale bars represent 500 µm (A-D), 20 µm (a-d) and 10 µm (á-d′).

These results indicate that the completion of the first meiotic wave at 20 dpp is contemporary with the formation of axonemes in both cilia and flagella. This temporal correlation, combined with previous data about the cytoplasmic continuity among spermatocytes (Figure 4), suggests the existence of a regulatory mechanism that functionally links the processes of ciliogenesis and flagellogenesis during testicular development.

### Proteomic Analysis Identifies Potential Regulatory Factors of Ciliogenesis and Flagellogenesis

Given the critical relationship between gene expression and physiological functions, and the observation that ciliated spermatocytes emerge alongside flagellated spermatids at 20 dpp, we then aimed to identify shared regulatory factors for axoneme formation during ciliogenesis and flagellogenesis. To achieve this, we conducted a comparative proteomic study using testicular extracts collected from multiple developmental stages: 8 dpp, 15 dpp, 19 dpp -before ciliogenesis-, 21 dpp – after ciliogenesis-, and adult.

Proteomic analysis was performed using tandem mass tag (TMT)-based quantitative proteomics, a method that enables high-throughput, simultaneous quantification of thousands of proteins. Across all samples, a total of 4304 proteins were detected (Supplementary Table 2). Quantitative data analysis was performed using PEAKS software, and all statistical analyses of the identified and quantified proteins were carried out in R for differential expression analysis. Differentially expressed proteins (DEPs) were defined as those showing a significant change in abundance between samples, with an adjusted p-value < 0.05. A total of 621 DEPs were identified (Supplementary Table 3).

Using data from all ages studied, we constructed a cluster heatmap of relative DEP abundance from prepubertal to adult testes (Figure 6-I and Supplementary Table 3). Gene Ontology (GO) analysis of DEPs revealed biological processes enriched during testicular development (Figure 6-II). Analysis identified four distinct DEP clusters. Two clusters showed significant abundance increase towards adulthood: proteins involved in sex reproduction and spermatozoa formation (cluster labelled in blue in Fig. 6-II) and those related to metabolic processes (cluster labelled in orange in Fig. 6-II). Remarkable proteins that increase their expression towards adulthood in the two abovementioned clusters are synaptonemal complex component SYCP3 and SYCE1 ^57^, lactate dehydrogenases A (LDHA) and C (LDHC, the testis-specific isoform) ^58^, metabolism regulators (HYOU1), and RNA/gene transcription regulators (PIWIL1, FXR1, SYDC). Interestingly, DCAF7 (the adaptor or substrate receptor in the CUL4-DDB1 E3 ubiquitin ligase complex) increases expression at 15dpp and peaks at 19dpp. Moreover, heat shock proteins and chaperones (HSP72, HSP90A, HSP74L, HSP71L), BiP (endoplasmic reticulum chaperone) and CAL (calreticulin) also showed strong increased expression towards adulthood (Fig. 6-II and Supplementary Table 3). These trends support the hypothesis that the testis undergoes a hypermetabolic phase as it develops and transitions towards tissue maturation. In contrast, another two clusters showed reduced expression with maturation: one involved in chromatin organization and localization, nucleosome assembly (green cluster in Fig. 6-II), and the other in RNA processing (black cluster in Fig. 6-II). This reduction is expected in a tissue that becomes functionally stable after development.

**Figure 6.**
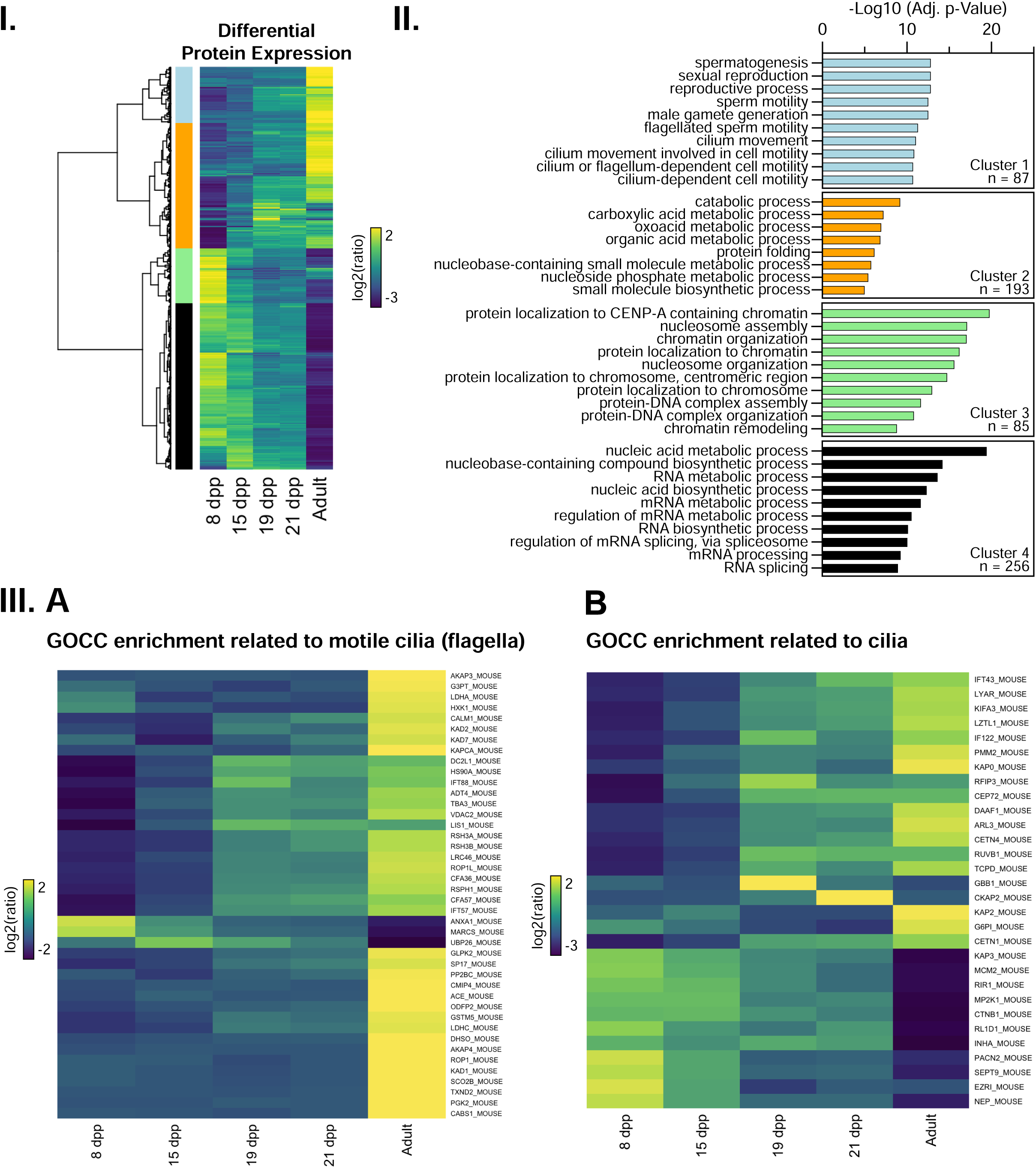
Comparative Proteomics Identifies Potential Regulators of Ciliogenesis and Flagellogenesis. **I.** Cluster heatmap of the relative abundance of the differentially expressed proteins (DEP) in testis from 8 dpp, 15 dpp, 19 dpp, 21 dpp, and adult animals. Data was clustered based on K-mean analysis. Differentially expressed proteins (DEPs) were defined as those showing a significant change in abundance between samples, with an adjusted p-value < 0.05. A total of 621 DEPs were identified and are shown in the heatmap. See Supplementary Table 2 for detailed data on each specific DEP. **II.** Gene Ontology (GO) enrichment analysis in the different biological processes for the four differentially expressed proteins clusters. **III.** Heatmap of dynamic abundances of the proteins associated with GO Cellular Component (GOCC) terms related to motile cilium (flagella) (**A**) and cilium (**B**).

To further explore cellular component associations (GOCC), we generated a heatmap of proteins related to axonemes of motile cilia, such as those in spermatids and spermatozoa. We observed a significant upregulation of several flagellar proteins. While LDHA is strongly expressed in adulthood, LDHC peaks at 19 dpp, supporting its known role in promoting flagellum formation ^58^. Additional proteins associated with flagellar assembly also showed increased expression (Fig. 6-III A), including heat shock proteins/chaperones (RSH3, HSPs, CALM) and the TRiC complex (TCPW, TCPE, TCPD, TCPA), which spiked from 15-19 dpp (Supplementary Table 3), coinciding with the age of emergence of ciliated spermatocytes and flagellated spermatids (Figure 5), suggesting that TRIC could be playing a role in axoneme assembly during this period. This data is consistent with increased expression from 15 dpp onwards of Tubulin-Binding Protein TBB3, while TBA8 peaks expression almost exclusive to adulthood. Other enriched proteins from 19 dpp onwards include Intraflagellar transport proteins (IFT57, IFT88), dynein complex-related proteins DC2L1 (DYNC2LI1) that plays a significant role in the function of flagella through its involvement in the dynein-2 complex and the IFT process ^59^; and Lis1 (Lissencephaly 1), which regulates microtubule dynamics during meiosis ^60^. In addition, AKAPs proteins, A-kinase anchoring proteins AKAP3 and AKAP4, known to localize in the flagellar fibrous sheath of the spermatozoa ^61^, were also increased toward adulthood. In contrast, proteins like ANXA1 (Annexin A1), which is involved in germ cell differentiation ^62^ showed higher expression in early stages of testis development (8 and 15 dpp).

We next created a specific GOCC heatmap of proteins linked exclusively to cilia (excluding those also found in motile cilia/flagella, therefore presumably corresponding to primary cilia although the category in the public GOCC database is labelled simply as ‘cilia’). Results showed that several proteins showed increased expression during testis maturation. Noteworthy examples are Intraflagellar transport proteins IFT22 and IFT43, which are enriched at 19 dpp, preceding the appearance of primary cilia in spermatocytes at 20dpp (Figure 2). ARL3 GTPase also became detectable at 19 dpp, aligning with related protein ARL13b presence in ciliated spermatocytes at 20 dpp (Figure 3). In addition, RFIP3 (RAB11FIP3, also known as FIP3) is detected from 19 dpp, in relation to its known implication in the Rab11-Rabin8-Rab8 ciliogenesis cascade, i.e., the trafficking of membrane proteins to the ciliary membrane ^63^. Moreover, centriolar proteins such as CETN1, CETN4, and centrosomal protein CEP72, a centriolar satellite component intricately involved in the formation and function of primary cilia ^64^, were also expressed from 19 dpp onwards. The detection of RUVBL1 from 19 dpp, an ATPase involved in the cytoplasmic pre-assembly of ciliary protein complexes before their transport to the cilium ^65^ is also remarkable. Conversely, some cilia-related proteins decreased in expression towards adulthood: for instance, Ezrin, which helps dock basal bodies during ciliogenesis ^66^, and SEPT9, which localizes along the primary cilium axoneme ^67^. This suggest that they might be implicated in cilia dynamics in cells of immature testis, as could be undifferentiated Sertoli cells, in which cilia have also been previously reported ^39^

These findings offer a comprehensive roadmap of potential regulatory targets for future research into testis development and its relationship to fertility, laying the groundwork for both descriptive and functional studies on meiotic ciliogenesis in the mouse model.

Interestingly, no significant changes were observed in two key indicators of Hedgehog (Hh) pathway activity: GLI1 protein levels and the processing of GLI3 into its repressor form, GLI3-R, both widely used as reliable readouts of Hh pathway activation ^68^. These findings were consistently supported by both proteomic analysis (Figure 6) and validation Western blot analysis, which confirmed the lack of upregulation in GLI1 and GLI3, as well as a stable ratio between full-length GLI3 and GLI3-R (Supplementary Fig. 5A,B). Notably, we also observed the absence of SMO (Smoothened, a transmembrane signal transducer in the Hh pathway ^69^) labelling in prepuberal 21 dpp ciliated spermatocytes (Supplementary Fig. 5 C). As a positive control to confirm the antibody’s specificity, SMO was detected in ciliated ASZ cells, a basal cell carcinoma line (BCC) known for active Hh signaling ^70,71^ (Supplementary Fig. 5 D). Together, these results suggest that the Hedgehog signalling pathway remains largely unaltered during the 19– 20 dpp window.

### Interference with meiotic cilia formation causes meiotic defects

After identifying the presence of ciliated spermatocytes throughout all stages of prepubertal prophase I, our focus shifted towards investigating whether the alteration of cilia assembly and disassembly may cause defects in meiosis or spermatogenesis. The prepuberal model is particularly well suited for these analyses, since the simultaneous appearance of the cilia at 20 dpp in different cells of the seminiferous epithelium represents a unique and synchronous system.

First, to inhibit cilia polymerization, we treated organotypic cultures of seminiferous tubules with chloral hydrate (CH), a compound known to remove primary cilia in somatic tissues ^72-74^. We treated seminiferous tubules from 19 dpp prepuberal animals for 24 hours, when ciliogenesis has not yet started (Figure 2). This means that after CH culture, spermatocytes should behave as 20 dpp meiosis without cilia. We selected a 24-hour treatment period due to a significant increase in cell death (evaluated by the presence of cleaved caspase 3) observed with longer exposures (Supplementary Figure 6-I A-C). Initial analysis focused on assessing the compound’s lethality and setting the best conditions for concentration and treatment duration. Two concentrations (40 µM and 100 µM) were tested. The results demonstrated extensive deciliation, with almost no prophase I spermatocytes retaining cilia, after treatment with 100 µM CH during 24 hours (Fig. 7-I A-D), confirming that CH is an efficient depolymerizing agent of cilia in mouse spermatocytes.

**Figure 7.**
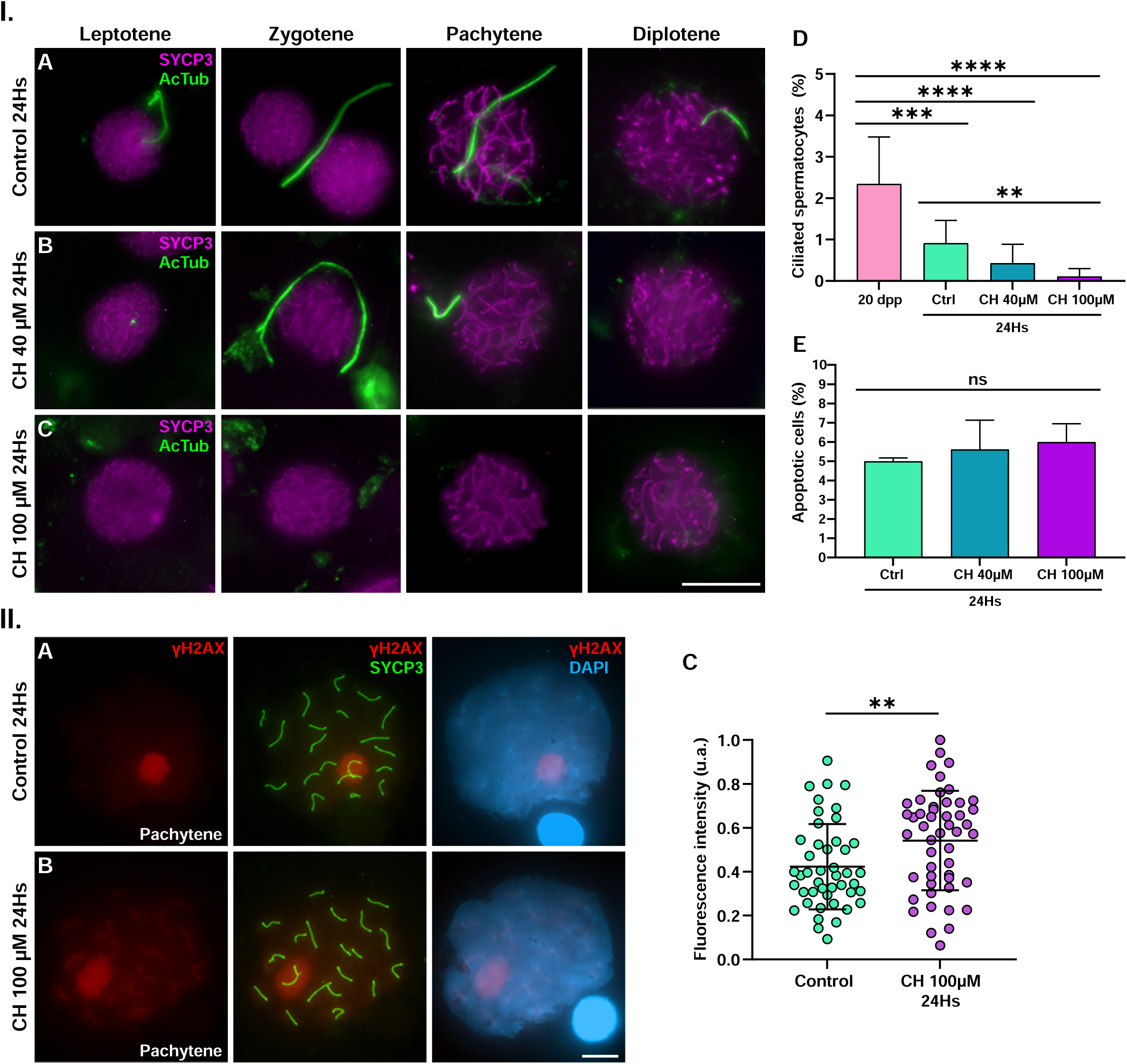
Deciliation induces persistence of DNA damage in meiosis. **I. Chloral hydrate (CH) causes complete deciliation in a 24-hour treatment in cultured spermatocytes. A-C.** Double immunolabelling of acetylated Tubulin (green) and synaptonemal complex protein 3 (SYCP3) (magenta) in cultured spermatocytes from 19 dpp mice. Experiments were conducted in control (A), 40µM HC (B) and 100µM HC (C) during 24 hours (24Hs). Images are shown for squashed spermatocytes in all stages of prophase I (leptotene, zygotene, pachytene and diplotene). All images are z-projection of 30-60 frames of stack images of each cell. Scale bar in C represents 10 µm. **D.** Graphical representation of the percentage of prophase I ciliated spermatocytes in fresh tissue from 20 dpp mice, compared to 24Hs cultured spermatocytes in control and CH-treated samples. Data represents mean ± SD, **** p < 0.0001, Χ2 test (a minimum of 1500 spermatocytes per condition, three biological replicates). E. Graphical representation of the percentage of apoptotic spermatocytes in 24Hs control and CH-treated cultured samples. Data represents no statistical significance (One-way ANOVA, p < 0.0001) (a minimum of 5500 spermatocytes per condition, three biological replicates). **II. A-B.** Double immunolabelling of SYCP3 (green) and DNA double-strand break marker γH2AX in spread mid-pachytene spermatocytes in 24Hs control (A) and 100µM HC (B) cultured samples. **C.** Dotplot of the fluorescence intensity (u.a.) of γH2AX signal in spermatocytes in 24Hs control and 100µM HC cultured samples. The increase in the fluorescence intensity is evident in treated spermatocytes. Data represents mean ± SD, **** p < 0.0001, Mann-Whitney test (n=45 spermatocytes per condition, three biological replicates).

To determine whether cilia disruption influences DNA damage repair, we assessed the persistence of DNA damage in pachytene spermatocytes following CH treatment. Quantification of apoptotic cells revealed no significant increase in cell death after 24-hours of treatment with either 40 µM or 100 µM CH (Fig. 7-I E), allowing us to rule out treatment-induced apoptosis as a confounding factor, at least for 24-hours. We subsequently analyzed mid-pachytene spermatocytes, assuming these cells were in early pachytene at the onset of treatment (Fig. 7-II A-B). Results showed that CH-treated cultures exhibited significantly higher levels of DNA damage compared to untreated controls, detected by marker of double strand breaks (DSBs) γH2AX (Fig. 7-II C), suggesting that disruption of meiotic cilia may impair DNA repair mechanisms during meiosis.

Second, to investigate the effects of blocking cilia depolymerization, we treated organotypic cultures of seminiferous tubules with MLN8237, a potent AURKA-specific inhibitor, at a concentration of 10 µM for 24 or 48 hours, following previously established protocols ^75^ (Fig. 8-I). We developed the experiment with cultured seminiferous tubules from 20 dpp mice (Figure 2). This means that after a 24-hours treatment, spermatocytes should behave as 21 dpp meiosis under Aurora A inhibition condition. No significant increase in apoptosis was observed after either treatment duration, as determined by cleaved caspase-3 staining in both control and MLN8237-treated spermatocytes (Supplementary Figure 6-II A-C). However, due to the progressive increase cell death, the 48-hours treatment was excluded from further analysis.

**Figure 8.**
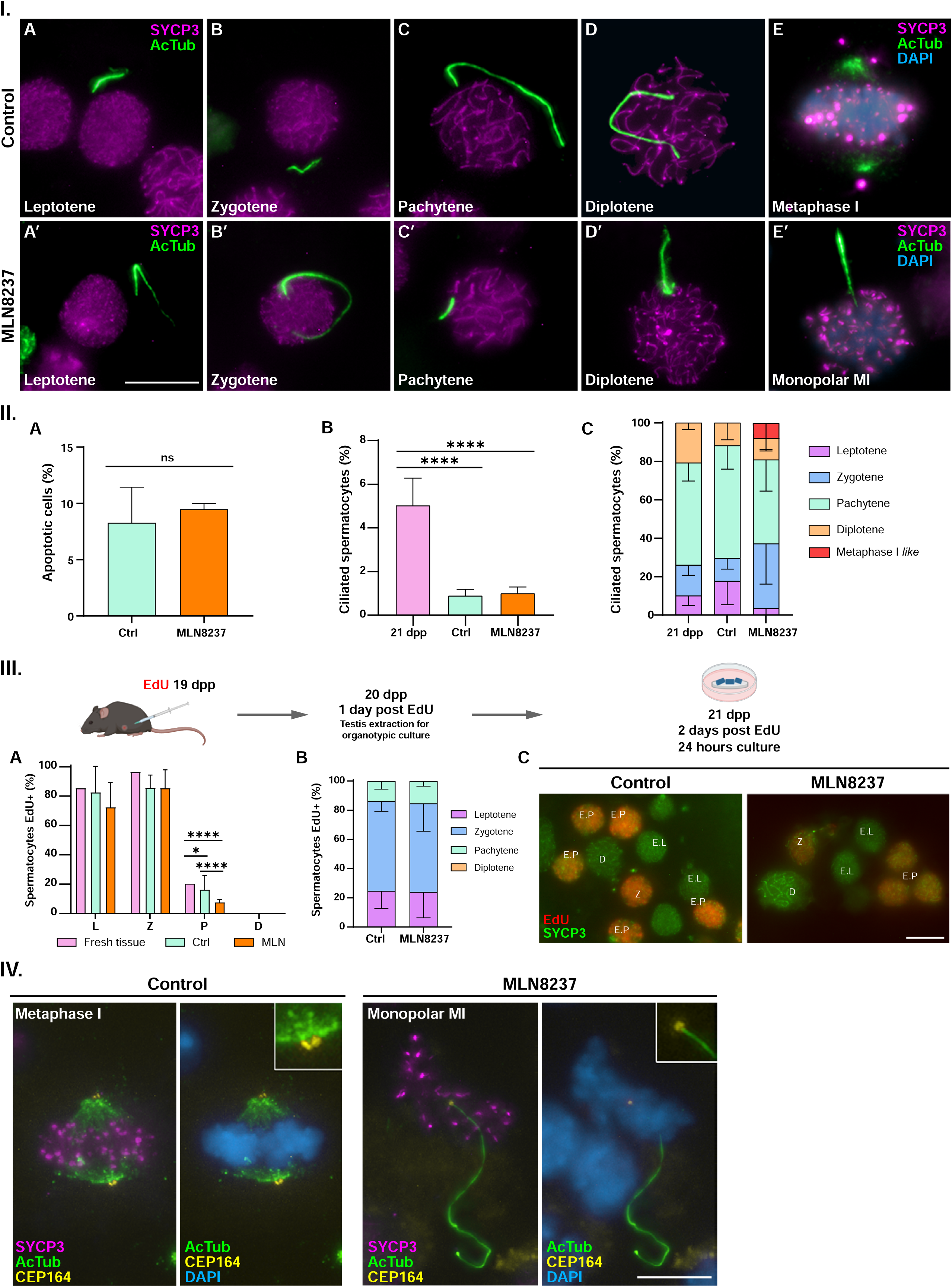
Aurora kinase A is a regulator of cilia disassembly in meiosis. **I. Aurora A inhibition causes interruption of cilia depolymerization. A-E.** Double immunolabelling of acetylated Tubulin (green) and synaptonemal complex protein 3 (SYCP3) (magenta) in cultured spermatocytes from 20 dpp mice. Experiments were conducted in 24-hours (24Hs) control and 10 µM MLN8237 73 cultured spermatocytes. Images are shown for squashed spermatocytes in all stages of prophase I (leptotene, zygotene, pachytene and diplotene) (A-D) and metaphase I (MI) (E) per each condition. All images are z-projection of 30-60 frames of stack images of each cell. Scale bar in A′ represents 10 µm. **II. Quantitative and qualitative analysis. A.** Graphical representation of the percentage of apoptotic spermatocytes in 24Hs control and MLN8237-treated cultured samples. Data represents no statistical significance (One-way ANOVA, p < 0.0001) (a minimum of 5500 spermatocytes per condition, three biological replicates). **B.** Graphical representation of the percentage of prophase I ciliated spermatocytes in fresh tissue from 21 dpp mice, compared to 24Hs cultured spermatocytes in control and MLN8237-treated samples. Data represents mean ± SD, **** p < 0.0001, Χ2 test, (a minimum of 1500 spermatocytes were quantified per condition, three biological replicates). **C.** Graphical representation of the distribution of ciliated spermatocytes in fresh tissue from 21 dpp mice, compared to cultured tissue for 24Hs in control and CH-treated cultured samples. **III. Edu pulse labelling indicates that Aurora A inhibition delays meiosis progression. A.** Graphical representation of the distribution of EdU-labelled spermatocytes among prophase I stages in control and 10 µM MLN8237 cultured samples. Χ2 test (a minimum of 300 spermatocytes were quantified per condition, three biological replicates). **B.** Image exemplification of cultured spermatocytes after EdU injection and organotypic culture. Double immunolabelling of EdU (red) and SYCP3 (green) in 24Hs control and MLN8237 cultured samples. E.L: early leptotene; Z: zygotene; EP: early pachytene; D: diplotene. Scale bar represents 10 µm. **IV. The inhibition of cilia depolymerization during late prophase I induces the appearance of monopolar ciliated metaphases I.** Triple immunolabelling of SYCP3 (magenta), acetylated Tubulin (green) and CEP164 in control bipolar metaphases I and MLN2537-treated monopolar ciliated metaphases I. All images are z-projection of 30-60 frames of stack images of each cell. Scale bars represent 10 µm.

Ciliated spermatocytes were observed in both control and MLN8237 24-hour treated samples (Fig. 8-I A–D and A′–D′). Quantitative analysis revealed that MLN8237 treatment did not significantly alter the overall number of ciliated spermatocytes between the control and the treated cultured samples (Fig. 8-II B). However, it did affect their distribution across meiotic stages, with a significant increase in ciliated spermatocytes observed during zygotene in treated samples (Fig. 8-II C). This could be due to a delay in cilia depolymerization or a delay in meiotic progression under AURKA inhibition. To distinguish between these possibilities, we performed EdU pulse labelling in 19 dpp mice, 24-hours prior to testis collection, enabling the tracking of meiotic progression (Supplementary Fig. 7-I). Analysis of histological cryosections and squashed spermatocytes confirmed the presence of ciliated spermatocytes in all Edu-labelled stages of prophase I (leptotene, zygotene, and pachytene) after 24, 48, and 72 hours post-EdU injection, consistent with previous findings in 20, 21, and 22 dpp mice (Fig. 2). Notably, diplotene spermatocytes were not labeled with EdU within this timeframe (up to 72 hours postinjection) (Supplementary Fig. 7-I A–F and 7-II), in line with the previously described duration of prophase I stages (Fig. 1). These results further confirm that EdU injection does not interfere with meiotic progression, as reported by ^76^. To determine whether MLN8237 treatment delays meiotic progression, we first confirmed that the proportion of EdU-labeled prophase I spermatocytes in treated cultures did not significantly differ from that in the fresh tissue or cultured controls (Supplementary Fig. 7-II A). We then compared the distribution of EdU-labeled leptotene, zygotene, and pachytene spermatocytes between control and treated cultures at the end of the experiment (24-hours post-culture and 48-hours post-injection). To ensure temporal consistency, we only analyzed EdU-labeled spermatocytes that had entered prophase I at the same time from now onwards. Our analysis revealed that AURKA inhibition does not lead to a delay in meiotic progression as the distribution of spermatocytes along the stages of prophase I does not change between the control and MLN-treated samples (Fig. 8-III A-B). Interestingly, although no significant differences were detected in the proportion of zygotene-stage spermatocytes in MLN8237-treated cultures compared to controls (Fig. 8-III A), a higher number of ciliated spermatocytes were observed under the Aurora A inhibition (Fig. 8-II C). These findings suggest that AURKA inhibition delays cilia depolymerization.

A striking finding of these experiments was the observation of ciliated spermatocytes at metaphase I. These spermatocytes were clearly anomalous and exhibited monopolar spindle phenotypes (Fig 8-IV). This is consistent with AURKA inhibition disrupting cilia disorganization ^28^, centrosome migration and spindle bipolarity ^75,77^. While bipolar spindles stained positively for AcTub, monopolar spindles lacked this signal. This phenotype closely resembled that observed following PLK1 inhibition with BI2536 ^75,78^, where AcTub is also absent in monopolar spindles. This was further validated by the absence of auto-phosphorylated AURKA (detected by AURKTph antibody as previously reported ^79^) in monopolar spindles treated with a 24-hours AURKA inhibition (Supplementary Fig. 8 A–C). In MLN8237-treated cultures, a further examination of CEP164, a marker of the mother centriole, revealed additional insights. In control metaphase I spermatocytes with bipolar spindles, four CEP164 signals were detected at opposite poles, aligning with our previous observations as well as those reported by other authors ^43,77^. However, in ciliated monopolar spindles from MLN8237-treated cells, CEP164 labeling was restricted to a single focus at the mother centriole, the one from which cilium emanates. This pattern suggests impaired distal appendage formation and defective centrosome migration (Fig. 8-IV). These results suggest that centrosome migration towards opposite poles is incompatible with the presence of centrosomes harbouring a polymerized ciliary axoneme. Alternatively, it implies that cilia disassembly might precede centrosome migration, suggesting the two processes are mutually exclusive. Together, these findings highlight AURKA as a key executor of cilia disassembly in mouse spermatocytes and emphasize its critical role in ensuring proper spindle assembly and meiotic progression.

## DISCUSSION

This study provides the first evidence of ciliated spermatocytes across mammalian species, offering novel significant insight into germ cell biology. Previous reports using electron microscopy identified primary cilia in somatic testicular cells, including peritubular myoid cells in rabbits ^80^, rats ^81^ and humans ^82,83^, and more recent work reported cilia in inmature Sertoli cells in the human testis^39^, and fetal Leydig cells in mouse ^84^. However, these studies did not document primary cilia in meiotic germ cells. Our previous work was the first to report ciliated spermatocytes in adult mouse testes ^43^ and the present findings now extend this observation to prepuberal mice and adult human spermatocytes.

This evolutionary conservation of primary cilia in meiotic cells suggests a previously overlooked but potentially fundamental role for these structures in coordinating meiotic progression, facilitating intercellular communication, or maintaining structural integrity within the germ cell cyst. Our findings open new avenues for understanding the regulatory functions of cilia in male fertility and raise important questions about their roles in testis development across mammals.

### Puberty: The foundation of fertility establishment in mouse

Our work advances the understanding of prepubertal spermatogenesis in mammals by identifying meiotic ciliogenesis as a defining event that distinguishes prepubertal from adult meiosis (Figure 9). In mouse, the first postnatal week is characterized by a distinctive program that lacks the self-renewing spermatogonia stage present in subsequent spermatogenic cycles. After 8 dpp, multiple overlapping spermatogenic waves arise from self-renewing undifferentiated spermatogonia ^49^, eventually culminating in the establishment of full fertility. Our study contributes to a deeper characterisation of mouse testis development by refining the developmental timeline: we identify early spermatids appearing in the lumen of seminiferous tubules at day 19, maturing into spermatids by day 28, and confirm that full epididymal maturation occurs at day 60, coinciding with previous reports ^85^.

**Figure 9.**
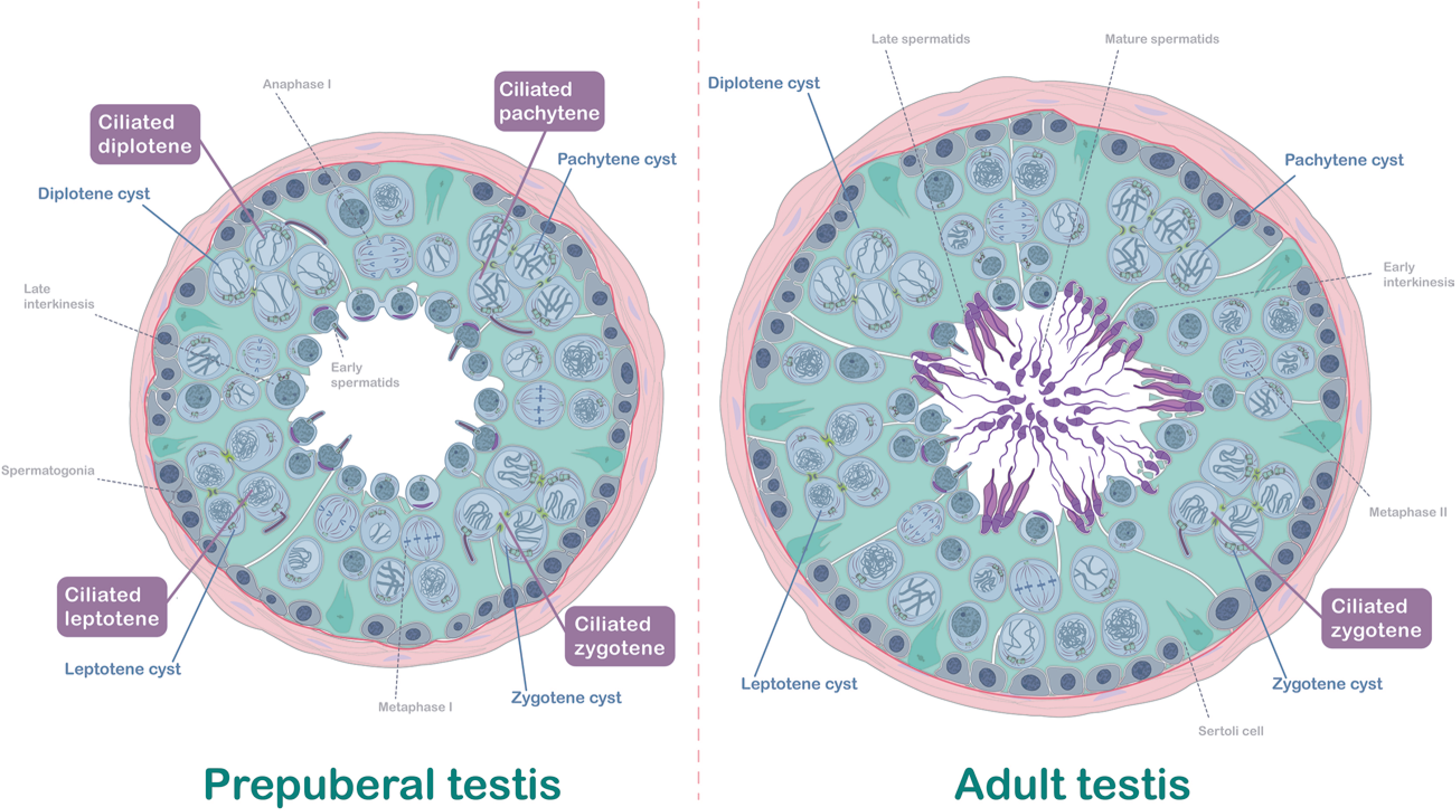
Schematic representation of the prepuberal versus adult seminiferous epithelium. Scheme represents a transversal section of a seminiferous tubule in testis of prepuberal and adult mouse. Spermatogonia (dark grey), spermatocytes (grey) and spermatids (purple) are embedded on the cytoplasm of Sertoli cells (turquoise). Centrosomes are indicated for every cell, in correspondence with timing of first centrosome duplication at zygotene, or second centrosome duplication at interkinesis. Cilia are represented in one of the prophase I spermatocytes of the cysts in prepuberal and adult mice. Intercellular bridges are indicated connecting the cytoplasms of the spermatocytes within the same cyst (highlighted in green). Myoid cells of the peritubular region are located surrounding the tubule (red).

Our use of the mouse model further reveals that spermatocyte ciliation begins during puberty, reinforcing the idea that this is a conserved and previously overlooked feature of mammalian spermatogenesis. Our work highlights the 19–20 dpp window as a pivotal stage, during which crucial events such as ciliogenesis and flagellogenesis are initiated. At the cellular level, we observed a small subset of spermatocytes exhibiting primary cilia during prophase I. Remarkably, these cilia are present across all substages of prophase I (leptotene, zygotene, pachytene, and diplotene), ruling out a direct correlation between cilia formation and the initiation of synapsis or desynapsis. This finding contrasts with zebrafish model, where development of cilia by zygotene spermatocytes has been linked to the initiation of synapsis and bouquet formation ^41^. However, consistent with our findings in prepubertal mice, other studies have observed cilia across all prophase I stages in zebrafish spermatocytes ^42^. In mouse, we found that ciliated spermatocytes appear only after 20 dpp, correlating with the completion of the first spermatogenic cycle and the formation of early spermatids. This likely explains why previous studies in spermatocytes using 12 and 16 dpp mice could only detect extremely short axonemal structures that barely extended the basal bodyu ^41^. Conversely, we detected extremely long cilia averaging 22 μm in length—substantially longer than primary cilia in other mouse tissues (e.g., 0.5–3.0 µm in brain, 1.5–3.0 µm in kidney proximal tubules, 5.0–7.0 µm in tracheal epithelium, ^86^). Given that it has been suggested that longer cilia enhance cellular responsiveness to extracellular signals in several tissues ^87,88^, we propose that the extremely long cilia observed in prepubertal spermatocytes may facilitate efficient intercellular communication within cysts during critical stages of testicular maturation.

Histologically, 20 dpp also marks the emergence of immature spermatid flagella at the tubule lumen. This synchronous appearance suggests a functional link between ciliogenesis and flagellogenesis during meiosis in prepubertal mice. To gain a broader understanding of the molecular mechanisms underlying these processes, we performed a global quantitative proteomic analysis of testis extract at different ages, prior or after meiotic first ciliogenesis. Previous studies addressed the different gene expression profile through spermatogenesis ^37,89-91^. In the context of prepuberal spermatogenesis, other authors identified differentially expressed proteins during meiotic initiation at 8 and 10 dpp ^48^, or the different expression pattern between 14 and 21 dpp ^92^. However, specific data for the 19-21 dpp window are absent.

We confirmed expected results in comparative proteomics in the 8-60dpp timeframe. For instance, proteins such as LDHC, crucial for glycolytic metabolism ^93^ and PIWIL1, a key piRNA pathway component maintaining genome integrity ^94^, showed progressive increases toward adulthood. Most interestingly, heat shock proteins such as HSP74L and HSP90A also showed robust upregulation, aligning with their established roles in maintaining spermatogenesis and sperm morphology ^95,96^. Our results also provided clear examples of progressive expression for HSP74L, whose absence causes infertility in the knockout mice due to altered sperm head morphology ^97^.

While the low abundance of ciliated spermatocytes suggested that identification of specific ciliogenesis regulators might be challenging, our analysis uncovered several promising candidates for regulators of Tubulin dynamics during meiosis. We observed upregulation of tubulin folding chaperones, particularly TRiC complex components (TCPG, TCPE, TCPD, TCPA), especially from 19dpp onwards. The TRiC complex is a key molecular chaperone complex facilitating the folding of tubulin, critical for ciliary axoneme and flagellar assembly. Accordingly, TRiC has been implicated in ciliogenesis via its interaction with the BBsome/IFT system ^98,99^. Notably, the timing of increased expression observed in our proteomic analysis aligns with the observation of both cilia and flagella at 20 dpp in the immunostaining cellular and histological analysis, suggesting that TRiC could be playing a role in axoneme assembly during this period. This data is consistent with increased expression from 15 dpp onwards of Tubulin-Binding Protein TBB3, a highly expressed protein in male germ cells ^100^ and identified by mass spectrometry in mammalian ciliary microtubules ^101,102^, which has been involved in microtubule dynamics required for both cilia and flagella formation ^103,104^. Expectedly, we detected high levels of TBA8 during adulthood, which is specific to spermatids with role in the mature stages of flagellar assembly and sperm motility ^3,105^.

Among other notable findings related to microtubule dynamics, SHCBP1L (Testicular Spindle-Associated Protein) was overexpressed at 19 dpp, consistent with its role maintaining spindle stability and cytokinesis during meiosis ^106^. In addition, we found an increased expression of DCAF7 in 19 dpp testis. DCAF7 is essential for male fertility ^107^ and islinked to ciliogenesis through interactions with DYRK1A/B in somatic cells ^108^. It is known to interact with DYRK1A and DYRK1B, which are kinases highly expressed in spermatids ^109^. Notably, DYRK1A is essential for microtubule assembly and ciliogenesis ^110,111^. It will be therefore exciting to address the interplay between DCAF7, DYRK1A, and tubulin isoforms such as TBB3 and TBB4B in future studies in spermatogenesis.

We also detected cilia-associated proteins such as ARL3, RFIP3, RUVBL1, and several IFT components (IFT57, IFT88, IFT22, IFT43), with protein levels rising at the same developmental window as ciliated spermatocytes’ appearance. The temporal concordance of the cytological expression of ARL13B and IFT88 further supports these findings. Finally, AKAP (A-kinase anchor proteins), known for their association with flagellar motility and structural integrity ^112^, were highly expressed in adulthood, coinciding with the completion of spermatogenesis and the establishment of full fertility.

In conclusion, our proteomic analysis not only validates existing knowledge but also provides new insights into the molecular events underlying ciliogenesis and flagellogenesis in the testis. Our data reveal previously uncharacterized proteins potentially involved in this process, bridging the gap between early developmental events and adult fertility. In this regard, Fang et al. recently offered a highly complete and complex comprehensive study using flow cytometry-isolated cells, where among other outstanding results, identified clear expression of cilium morphogenesis and cilium assembly proteins in mid-pachytene spermatocytes compared to round spermatids ^113^. These studies align with our data and collectively suggest a compelling link between ciliogenesis and testis development. Future research should aim to dissect the molecular pathways regulating ciliogenesis during male meiosis, offering deeper insight into its role in testicular maturation and long-term fertility maintenance.

### Meiosis cilia share common regulators with somatic cells in mouse

Our findings suggest shared regulatory mechanisms governing meiotic and somatic cilia dynamics. The use of CH is a strategy previously used to induce deciliation in somatic cells, as it chemically disrupt cytoskeletal integrity by destabilizing microtubules and ciliary anchorage to the basal body ^72-74^. This study is the first to employ CH treatment in cultured spermatocytes. Our results demonstrate that a 24-hours CH exposure effectively induced complete deciliation in prepubertal spermatocytes, without causing significant cell death. Moreover, CH does not induce genotoxic effects in previous reports ^114^, although it has been reported to trigger the generation of ROS (Reactive Oxygen Species) at higher concentrations ^115^. Interestingly, primary cilia play a vital role in regulating autophagy and DNA damage repair in glioblastoma cells ^107^. In our study, CH treated spermatocytes exhibited higher levels of γH2AX, which could be caused by an impairment in DNA repair during prophase I, suggesting a link between cilia integrity and efficient DSBs repair. This observation aligns with findings in zebrafish, where cilia depletion delays DSB repair, impairs crossover formation, and increases apoptosis, ultimately delaying— but not arresting—meiotic progression ^42^. Together, these findings suggest a conserved role for cilia in facilitating DNA repair during meiosis across vertebrates. In somatic cells, the presence of DNA Damage Response (DDR) proteins, such as BRCA1, ATM, ATR and CHK1, has been reported at the centrosome ^116^. This suggests a potential link between primary cilia, centrosome and DNA repair during meiosis. Future studies are needed to unravel the precise molecular mechanisms connecting cilia maintenance to DSB resolution in male mouse meiosis.

We also examined the inhibition of cilia depolymerization in cultured spermatocytes. Aurora kinase A (AURKA), a master regulator of centrosomal functions during mitosis ^27,117,118^, is known to trigger cilia disassembly prior to cell division ^28,29^. Notably, AURKA also interacts with centriolar IFT88 ^119^. Moreover, IFT88 has also been detected at somatic centrosomes and is required for spindle orientation in mitosis ^120^. In the present work, we have described IFT88 located at centrioles. Accordingly, previous studies have also detected AURKA at meiotic centrosomes ^75,79,121^. This suggests a functional interaction between AURKA and IFT88 in spermatocytes and open new research avenues linking ciliary proteins, centrosome dynamics and spindle organization.

Microtubule remodelling, essential for centrosome migration, is modulated by the acetylation status of tubulin. Tubulin deacetylation enhances microtubule turnover and flexibility, facilitating reorganization required for spindle bipolarity. Conversely, tubulin acetylation stabilizes microtubules, reducing their propensity for depolymerization and impeding centrosome migration. In somatic cells, cytoplasmic deacetylases HDAC6 (Histone Deacetylase 6) ^122,123^ and Sirtuin 2 ^124^. mediate tubulin deacetylation, whereas α-tubulin acetyltransferase 1 (ATAT1) promotes microtubule stabilization. ATAT1 depletion impairs Polo-like-Kinase 1 (PLK1) recruitment to centrosomes, affecting microtubule nucleation ^125^. While the roles of AURKA, HDAC6, and ATAT1 in tubulin acetylation/deacetylation and microtubule dynamics during mitosis are well-documented, their functions during spermatogenesis remain unclear. Notably, PLK1, AURKA ^75^ and IFT88 (this work) localize to meiotic centrioles, suggesting a conserved regulatory centrosome-cilium crosstalk during meiosis. In addition, HDAC6 ^126^ and ATAT1 had been pointed to play functional roles in mouse flagella ^127^, yet its potential regulatory interactions with AURKA remain to be elucidated in the context of meiosis cilia regulation.

Given AURKA’s multiple cellular pools with distinct substrate specificities ^128^, our study suggests that AURKA regulates the transition from ciliated to deciliated states during late prophase I in prepubertal meiosis. This role adds to AURKA’s function in centrosome migration ^75,77^. In somatic cells, cilia disassembly is necessary to release the basal body and enable spindle pole formation ^129^. Contrariwise, during meiosis, meiotic ciliogenesis begins once the first meiotic division is established -^43^ and this study-, therefore the real mutually exclusive events are cilia persistence and centrosome migration. Accordingly, cilia are never observed in control cells beyond early diakinesis, when centrosome migration begins ^75^, unless AURKA activity is inhibited, allowing cilia to persist up to metaphase I.

Finally, the presence of ciliated spermatocytes both before and after centriole duplication at zygotene raises important questions about the interplay between ciliogenesis and centrosome duplication. In somatic cells, PLK4 maintains centriolar satellite integrity and promotes ciliogenesis through PCM phosphorylation ^130^, but is inactive during G1 to permit cilia formation ^131^. In contrast, meiosis presents a unique scenario where PLK4 remains active throughout prophase I, driving centriole duplication ^132^. The concurrent kinase activity of AURKA, PLK1 and PLK4 during male meiosis ^75,77,132^ suggests a potential crosstalk, possibly through PCM phosphorylation or other mechanisms, facilitating an integrated regulatory network coordinating centrosome function and ciliary dynamics. The presence of ciliated spermatocytes throughout all stages of prophase I raises the possibility that PLK4, PLK1 and AURKA may cooperate to regulate ciliary assembly or disassembly, either before or after centriole duplication. Future investigations should focus on dissecting the functional interplay between these kinases, offering critical insights into the regulation of cilia in male meiosis and fertility.

## MATERIAL AND METHODS

### Materials

Testes from prepuberal and adult C57BL/6 (wild-type, WT) male mice were used for this study. Lactating prepuberal individuals were sampled at 8 to 30 days *postpartum* (dpp). Non lactating individuals of 30 to 60 dpp were also used. All animal procedures were approved by local and regional ethics committees (UAM Ethics Committee for Research and Animal Welfare) and performed according to European Union guidelines.

Human samples were provided by Dr. Ignasi Roig (Universidad Autónoma de Barcelona, Spain). All human samples donation were approved by the corresponding institution (Hospital de Bellvitge, Barcelona) with the ethical committee reference BB20-024. Samples used for this study correspond to references B03-8544F y la B05-20957J of the correspondent biobank.

### Squashing and spreading methodology and immunofluorescence microscopy

Seminiferous tubules were fixed and processed following previously described protocols for the squashing technique ^133^. For spreading technique, we also followed previous protocols ^134^.

Acetylated Tubulin was detected with a mouse monoclonal antibody (Sigma, T7461) at a 1:1000 dilution. SYCP3 was detected with a mouse monoclonal antibody (Santa Cruz, sc-74569) at a 1:200 dilution. SYCP1 was detected with a mouse monoclonal (Abcam, ab15090) at a 1:100 dilution. DSBs were detected with a rabbit monoclonal against histone H2AX phosphorylated at serine 139 (γH2AX) (Abcam, ab2893) at 1:500; CEP164 was detected with a rabbit polyclonal antibody (Sigma, ABE2621) at a 1:100 dilution; IFT88 was detected with a rabbit polyclonal antibody (Proteintech, 13967-1-AP) at a 1:100 dilution; TEX14 was detected with a rabbit polyclonal antibody (Proteintech, 18351-1-AP) at a 1:500 dilution; VASA was detected with a rabbit polyclonal against mouse DDX4/MVH (Abcam, ab13840) at a 1:100 dilution. Pools of phosphorylated Aurora kinase A/B/C (AURKTph) were revealed with a rabbit polyclonal anti-AuroraTph antibody (Cell Signaling, 2914S) at a 1:30 dilution. SMO was revealed with a rabbit polyclonal antibody (Abcam Ref.72130) at 1:400 dilution. Corresponding secondary antibodies were used against rabbit and mouse IgGs conjugated with either AMCA, Alexa 488, Cy3 or Alexa 647 (Jackson Laboratories), all of them used at a 1:100 dilution.

Immunofluorescence images and stacks were collected on an Olympus BX61 microscope equipped with epifluorescence optics, a motorized Z-drive, and Olympus DP74 digital camera controlled by Cellsens software (Olympus Life Science). Finally, images were processed with ImageJ (National Institute of Health, USA; http://rsb.info.nih.gov/ij) or/and Adobe Photoshop softwares.

### Histology

For histological cryosections, dissected testis were fixated with 4% PFA for 24 h, then they were inmersed in 15% and 30% sucrose solution 24 hours for each step. Last, they were included in OCT (Sakura Finetek Europe) and immediately frozen. The OCT blocks were cut in 10 μm thick sections using a Leica CM1520 cryostat. Slides were washed in PBS three timesfollowed by permeabilization with PBS/0.1% Triton X-100 for 20 minutes. Then an antigen retrieval protocol was performed with sodium citrate 0,1M pH 6,0 at 85°C for 30 minutes. After 3 washes in PBS, slides were blocked with PBS/2,5% BSA 0,1% Tween 20 for 30 minutes before being used for immunofluorescence protocol.

For human sections, samples were fixed in Formalin and included in paraffin. The paraffin blocks were cut in 7 μm thick sections. Then slides were dewaxed by washing three times in xylene, then in 100%, 96% and 75% ethanol two times per step, and finally in H_2_O. Then an antigen retrieval protocol was performed with sodium citrate 10mM, 0,5% Tween-20 pH 6,0 at 95°C for 20 minutes. After 2 washes in PBS-T, slides were permeabilized with PBS/0.1% Triton X-100 for 20 minutes. Finally, before being used for immunofluorescence protocol, slides were blocked for 1 hour using 0,3% BSA, 1% goat serum, glycine 0.00034M, 0,02% Triton X-100, 0,01% Tween-20 in PBS.

### Cell lines and culture and immunofluorescence microscopy

We used Basal Cell carcinomas (BCC) from the line ASZ, generously provided by Dr. Elisa Carrasco (Universidad Autónoma de Madrid), as positive control for the detection of active Hedgehog (Hh) pathway. For immunofluorescence protocol, cells were grown on coverslips were fixed in 3.7% formaldehyde/phosphate-buffered saline (PBS), permeabilized with 0.1% Triton X-100/PBS (1 hour at room temperature, RT) and incubated for 1 hour and 30 minutes at RT with the primary antibodies. anti-acetylated tubulin (1:1000; mouse; Sigma (Ref. T7451) and SMO (1:400; rabbit; Abcam Ref.72130). Corresponding secondary antibodies were used against rabbit and mouse IgGs conjugated with either Alexa 488 or 549 (Jackson Laboratories) at a 1:200 dilution, incubated for 1 hour at RT. Finally, cells were counterstained with Hoechst for 5 minutes and mounted in ProLong (Invitrogen). Images were taken using an epifluorescence microscope linked to an Olympus DP50 digital camera.

### Electron microscopy

Testis from 21 dpp prepuberal mice were fixed in a mixture of 4% PFA, 2% glutaraldehyde in 0.1 M sodium cacodylate buffer, for 24 hours. After fixing, samples were postfixed in a mixture of 1% osmium tetroxide and 0.8% potassium ferrocyanide, for an hour. Samples were treated with uranyl acetate 2% *en bloc* staining and dehydrated through a graded series of ethanol (30%, 50%, 70%, 95% and 100%), then embbebed in an acetone – Durcupan sequence and transferred to Durcupan epoxy resin. Ultrathin sections of 60 nm were obtained with Leica ultramicrotome and placed on grids. Finally, sections were stained with 2% uranyl acetate and Reynolds lead. For ultrastructural study, we used a Transmission Electron Microscope Jeol Jem 1010 (Tokyo, Japan) equipped with a Gatan Orius SC200 (Pleasenton, Ca) digital camera.

### Organotypic culture of seminiferous tubules and inhibition treatments

Culture of seminiferous tubules was performed as previously described ^75,135^, following previous reviewed recommendation ^136^. Testes from WT prepuberal mice were removed, detunicated and fragments of seminiferous tubules were cultured at 34°C in an atmosphere with 5% CO_2_ onto agarose gel half-soaked in MEMα culture medium (Gibco A10490-01) supplemented with KnockOut Serum Replacement (KRS) (Gibco 10828-010) and antibiotics (Penicillin/Streptomycin; Biochrom AG, A2213). Organotypic cultures were treated with 40-100 µM of aqueous chloral hydrate (CH) (Sigma-Aldrich, C8383) for 24 or 48 hours to inhibit primary cilia polymerization. To inhibit AURKA, 10 µM of MLN8237 – also called Alisertib – (Selleckchem, S1133) diluted in 10% DMSO was added to the culture medium and seminiferous tubules were also recovered after 24 or 48 hours of treatment. Controls for each experiment were done with seminiferous tubules cultures in MEMα culture medium and 10% DMSO. After 24 or 48 hours, control and inhibitor-treated seminiferous tubules were subjected to the squashing or spreading technique (described above). To inhibit PLK1, 100 µM of BI2536 was administrated during 24 hours to organotypic culture of seminiferous tubules as previously reported ^75^.

### EdU pulse labelling

19 dpp prepuberal mice were injected intraperitoneally with 30 μl of 10 mM 5-ethynyl-2-deoxyuridine (EdU) (Thermo Fisher Scientific) diluted in PBS, and sacrificed after 24 hours, 48 hours, or 72 hours, a timeframe that corresponds to 20 dpp, 21 dpp and 22 dpp. This allowed us to cover the ages when ciliated spermatocytes first appear at the testicular tissue tracking the first meiotic wave. Testicular samples were processed as described above and spermatocytes were labelled with anti-Acetilated Tubulin and anti-SYCP3. EdU was revealed using the Click-iT labelling as recommended by the supplier. For organotypic culture experiments, testis were extracted 24-hours after EdU injection and cultured for 24 additional hours and processed as described in previous section.

### Quantification and Statistical analysis

All values included in the figures are detailed in the supplementary material (Supplementary Table 1).

For the description of the first meiotic wave in prepuberal mice 2000 spermatocytes were quantified in two biological replicated for each age (between 8 to 60 dpp).

For the quantitative analysis of the presence of ciliated spermatocytes, 500 spermatocytes in prophase I were analyzed in three biological replicates per condition (between 20 to 60 dpp). The length of cilia was quantified in a minimum of 30 spermatocytes per stage, for each analyzed age condition (20 dpp, 21 dpp or 22 dpp).

For the organotypic cultures experiments using CH or MLN8237, 500 spermatocytes in prophase I were analyzed in three biological replicates per condition (control, CH and MLN8237).

For the quantification of DNA damage marked by γ-H2AX, quantification of the immunofluorescence intensity was estimated by measuring the integrated fluorescence density in individual nuclei using ImageJ by creating a binary mask with the DAPI staining. Then the CTCF (Correlated Total Cell Fluorescence) was calculated. The acquisition time was fixed for all acquired images, and the quantification was only performed using the original unmodified images. A minimum of 15 pachytene spermatocytes per condition were analyzed in each experiment, with three biological replicates.

All statistical analyses were performed using GraphPad Prism 10 (GraphPad Software Inc.). For the analyses of differences in the number of cilia present a Χ^2^ test was performed. For the rest of the analyses, first, a normality test was performed for each analysis. Differences were analyzed by Mann-Whitney or one-way ANOVA when the data followed a normal distribution, and by a Kruskal-Wallis test when they did not. Then, a multiple comparison test was performed. Values are expressed as mean±SD and P-values below 0.05 (P < 0.05) were considered statistically significant.

### Proteomic Informatics Analysis

The testes were decapsulated and homogenised with manual grinder in RIPA buffer (Sigma) supplemented with phosphatase and protease inhibitor cocktails. Following, the lysates were centrifuged at 12000 g for 15 min to remove insoluble material. Protein concentration was determined using the Pierce BCA Protein assay, and samples were separated by SDS-PAGE to assess protein quality. Subsequently, proteins were concentrated by running them on a stacking SDS-PAGE gel and digested using trypsin. The resulting tryptic peptides were desalted, concentrated and labelled with TMT 6-plex reagents following the manufacturer’s instructions. The efficiency of TMT labelling was evaluated through a preliminary LC-MS/MS analysis, and data quality was assessed accordingly. The TMT-mix was then fractionated using reverse-phase spin columns at basic pH. Each of the resulting fractions was analysed by LC-MS/MS on an LTQ-Orbitrap Velos Pro mass spectrometer using a short gradient method.

Quantitative data analysis was performed in PEAKS software and all statistical analysis of the identified and quantified proteins were performed in R statistical software (http://www.R-project.org/, version v4.4.2), DAVIS and GraphPad software (Prism 10, version 10.4.2). Tandem mass spectra were searched against the Swiss-Prot mouse database, applying a 1% false discovery rate (FDR) at both the peptide and protein levels. Proteins were considered confidently identified when supported by at least two unique peptides, with a protein-level FDR below 0.01. The relative abundance of each protein was defined as the log2 of the ratio between its abundance at a specific testis maturation stage and its mean abundance across all testis maturation stage Differential Expressed Proteins (DEPs) among the five stages of testis maturation were defined by two-way ANOVA filtered with an adjusted q value <0.05. To identify DEP groups with similar abundance patterns during different stages of testis maturation, the optimal number of clusters (k value) was first determined using the Cluster package in R (version 4.3.3). Subsequently, relative abundance values of the DEPs were clustered using K-means analysis. DEPs were subjected to Gene Ontology (GO) enrichment analyses using the DAVID version v2024q4 ^137,138^. Functional annotation clustering was performed with default settings, and enriched GO terms were considered significant upon an adjusted p-value (Bonferroni corrected) less than 10e-5.

### Western blot

Lysates were then processed for SDS-PAGE and Western blot as previously described (Barbeito et al. 2023 Frontiers Cell Dev Biol 11: 1190258). Briefly, 30 µg of each lysate were loaded into 8% SDS-PAGE gels. After transfer to nitrocellulose, membranes were blocked and incubated with the following primary antibodies: mouse anti-GLI1 (Proteintech, 66905-1-Ig, 1:1000), goat anti-GLI3 (R&D Systems, AF3690, 1:1000), mouse anti-GAPDH (Proteintech, 60004-1-Ig, 1:100,000), or mouse anti-alpha-tubulin (Proteintech, 66031-1-Ig, 1:10,000). Membranes were then washed and incubated with these secondary antibodies: horseradish peroxidase (HRP)-conjugated goat anti-mouse IgG (Thermofisher, A16072) or HRP-conjugated rabbit anti-goat IgG (Thermofisher, #31402). After three more washes, signal detection was performed using SuperSignal West Pico PLUS Chemiluminescent Substrate (Thermofisher, 34578) and a Fusion Solo S imaging system (Vilber).

## ACKOWLEDGEMENTS and FUNDING

We express our sincere thanks to Dr. Elisa Carrasco and Paula Alcaraz Laso (Universidad Autónoma de Madrid) for generously providing the basal cell carcinoma cultured cells for Hh positive control. We also thank Dr. Arántzazu Rodríguez de Gortázar and Dr. Juan Ardura for their valuable advice on the CH experiments. We thank Manuel Guerrero for preparing the scientific illustrations, and Natalia Calvo for her technical assistance.

This work was funded by BIOUAM05-2022 to R.G. and J.P., MCIN/AEI grants PID2022-140364NB-I00 to R.G. and J.P., PID2023-149472NB-I00 to F.G.G, and grants PID2022-138905OB-I00 and a grant from La Funació Marató de TV3 (677/U/2021) to I.R.

## AUTHOR CONTRIBUTIONS

RG, JP, IPM and PL conceived the study; IPM, PL, HZ, SPM, PB and MLP performed all experiments; all authors analyzed the results; RG, JP, IPM, PL, IR and FGG wrote the paper with some contributions from the other co-authors.

---

**Supplementary Figure 1:**
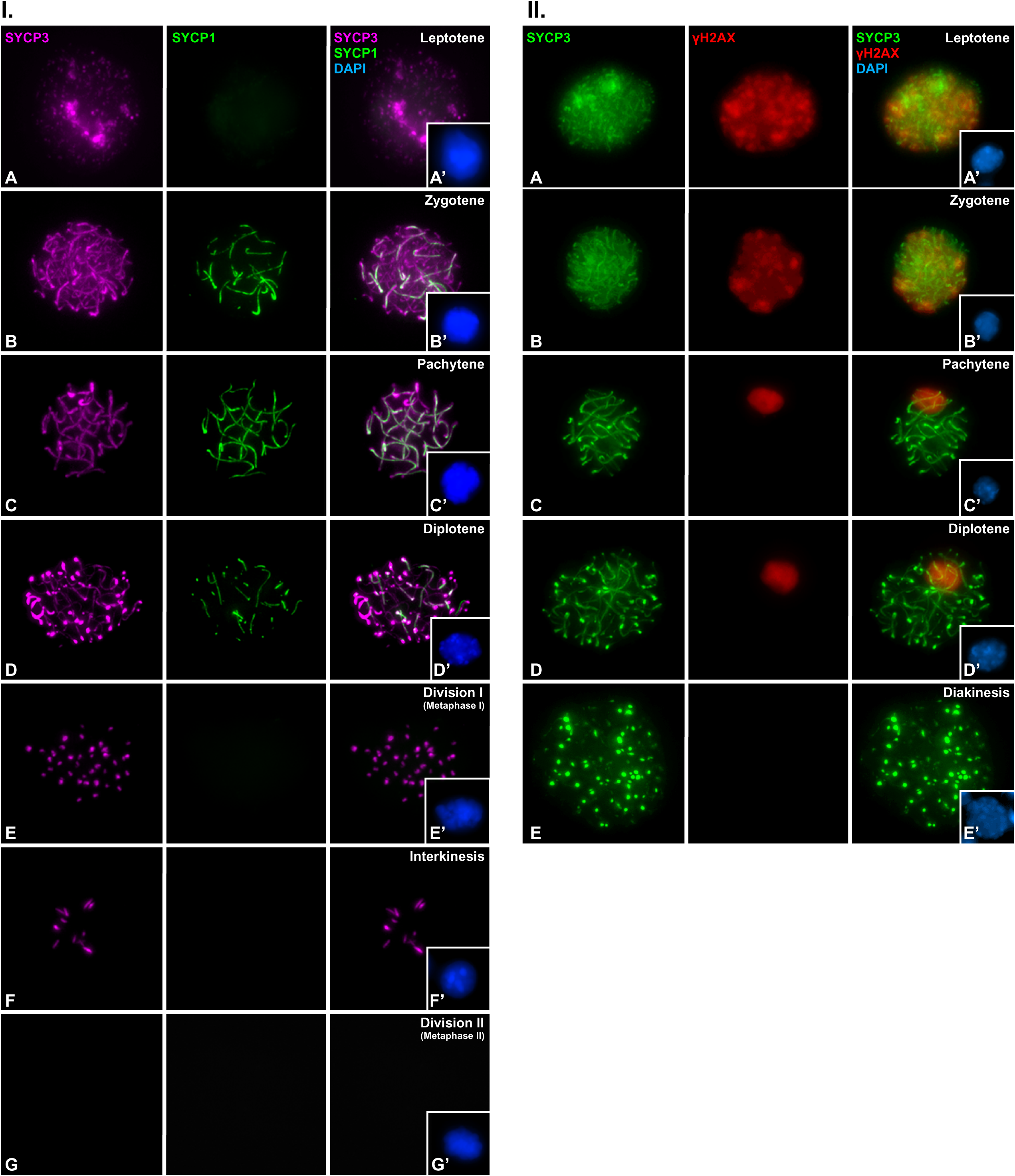
Meiotic stages during the first meiotic wave. **I. A-G.** Double immunolabelling of synaptonemal complex protein 3 (SYCP3) (magenta) and synaptonemal complex protein 1 (SYCP1). Images are shown for squashed spermatocytes in all stages of prophase I (leptotene, zygotene, pachytene and diplotene) (A-D), metaphase I (E), interkinesis (F) and metaphase II (G). All images are z-projection of 30-60 frames of stack images of each cell. Chromatin is stained with DAPI (blue) in A’-G’. Scale bar represents 10 µm. **II. A-E.** Double immunolabelling of synaptonemal complex protein 3 (SYCP3) (green) and DNA double-strand break marker γH2AX (red). Images are shown for squashed spermatocytes in all stages of prophase I (leptotene, zygotene, pachytene and diplotene) (A-D) and diakinesis (E). All images are z-projection of 30-60 frames of stack images of each cell. Chromatin is stained with DAPI (blue) in A’-E’. Scale bar represents 10 µm.

**Supplementary Figure 2:**
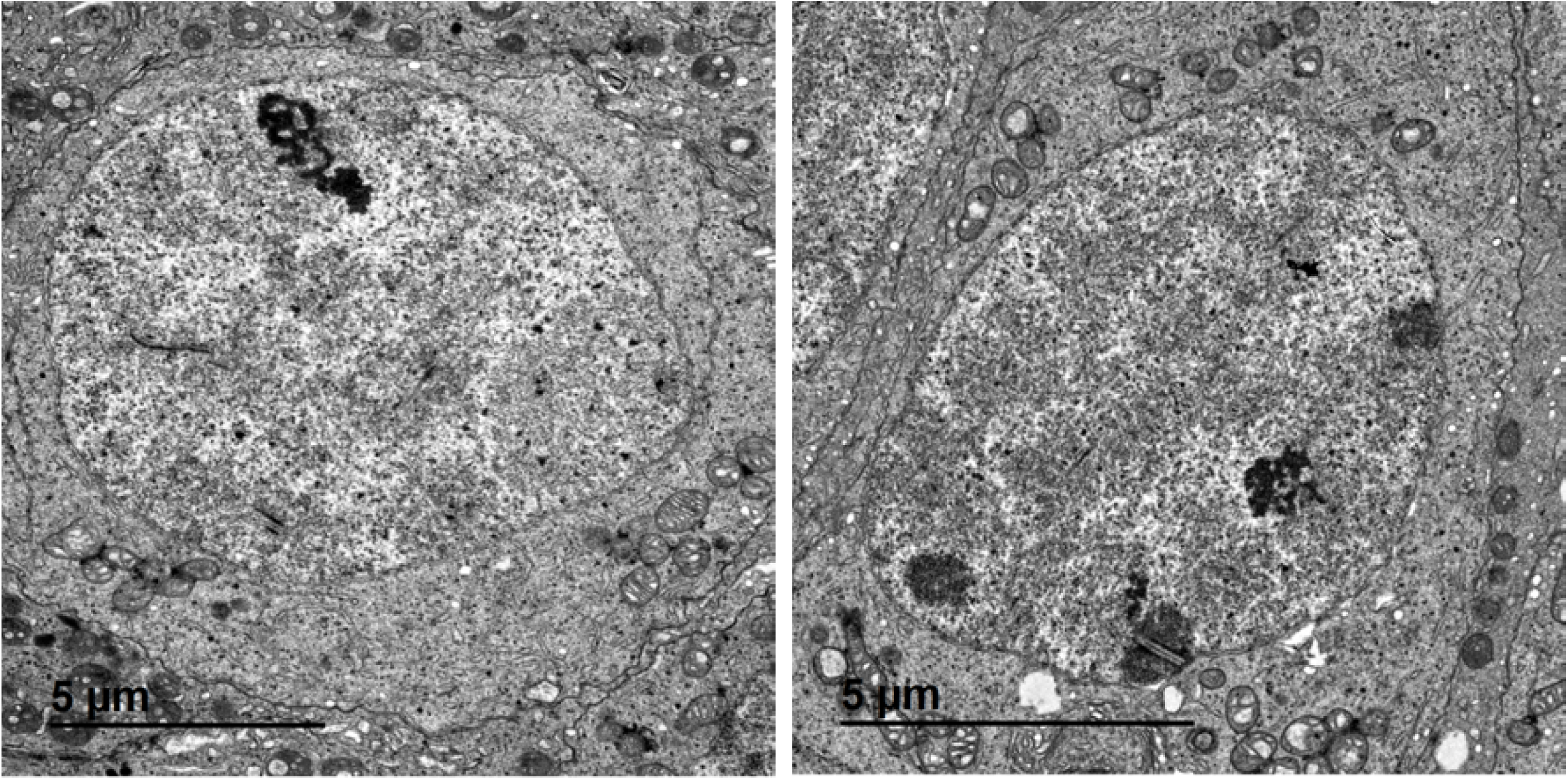
Ultrastructure of prophase I spermatocytes. Electron microscopy images of spermatocytes at prophase I presenting fragments of synaptonemal complex, and a prominent nucleolus. Spermatocytes are connected by intercellular bridges.

**Supplementary Figure 3:**
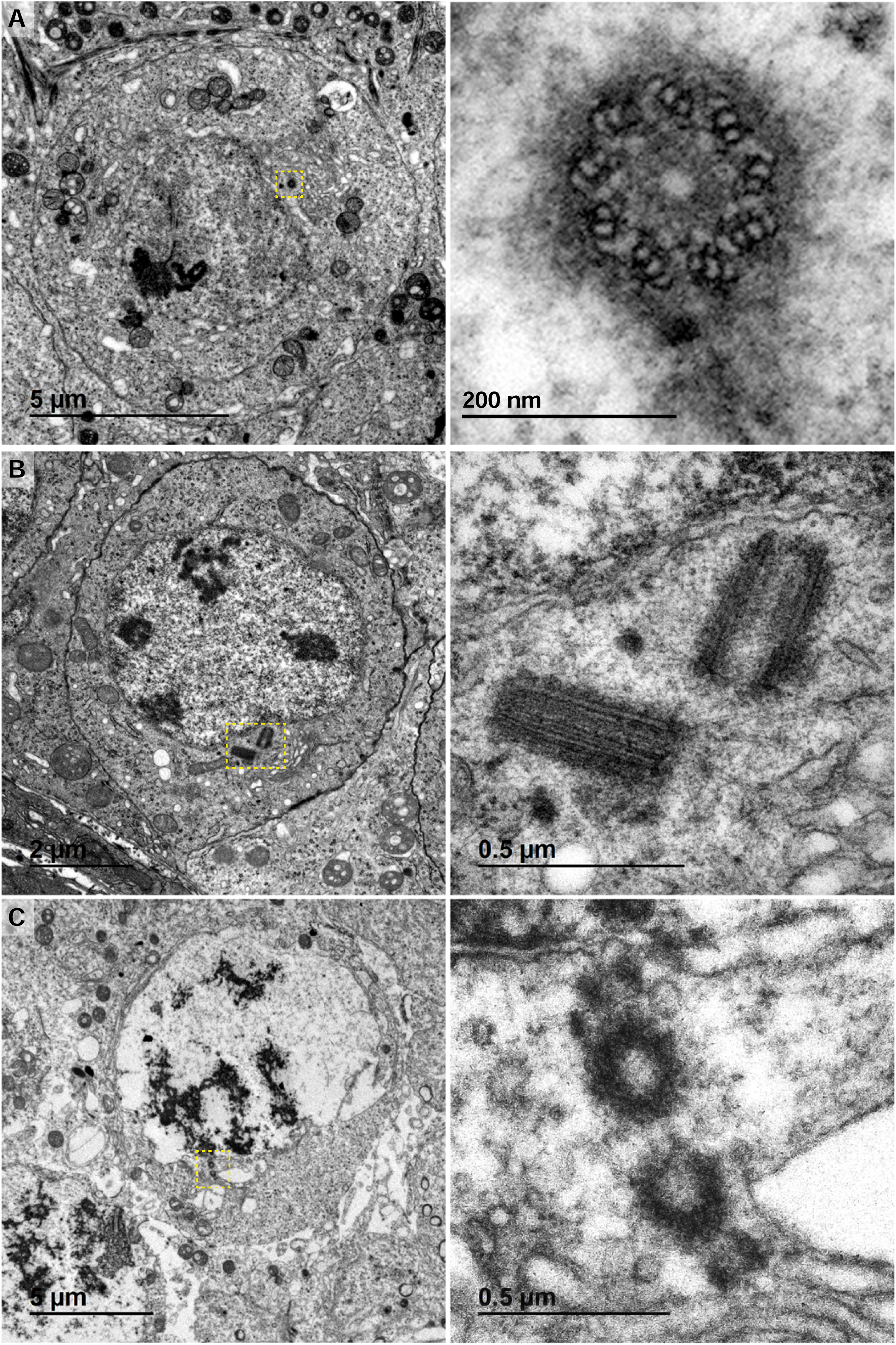
Meiotic centrosome under the electron microscope. **A-C**. Electron microscopy images of spermatocytes in early prophase prior to centrosome duplications (A and B), and after centriole duplication (C). Centrosomes are enhanced in the correspondent image. The pair of centrioles with their procentrioles are clearly seen during centrosome duplication (C). Scale bars are included in each image.

**Supplementary Figure 4:**
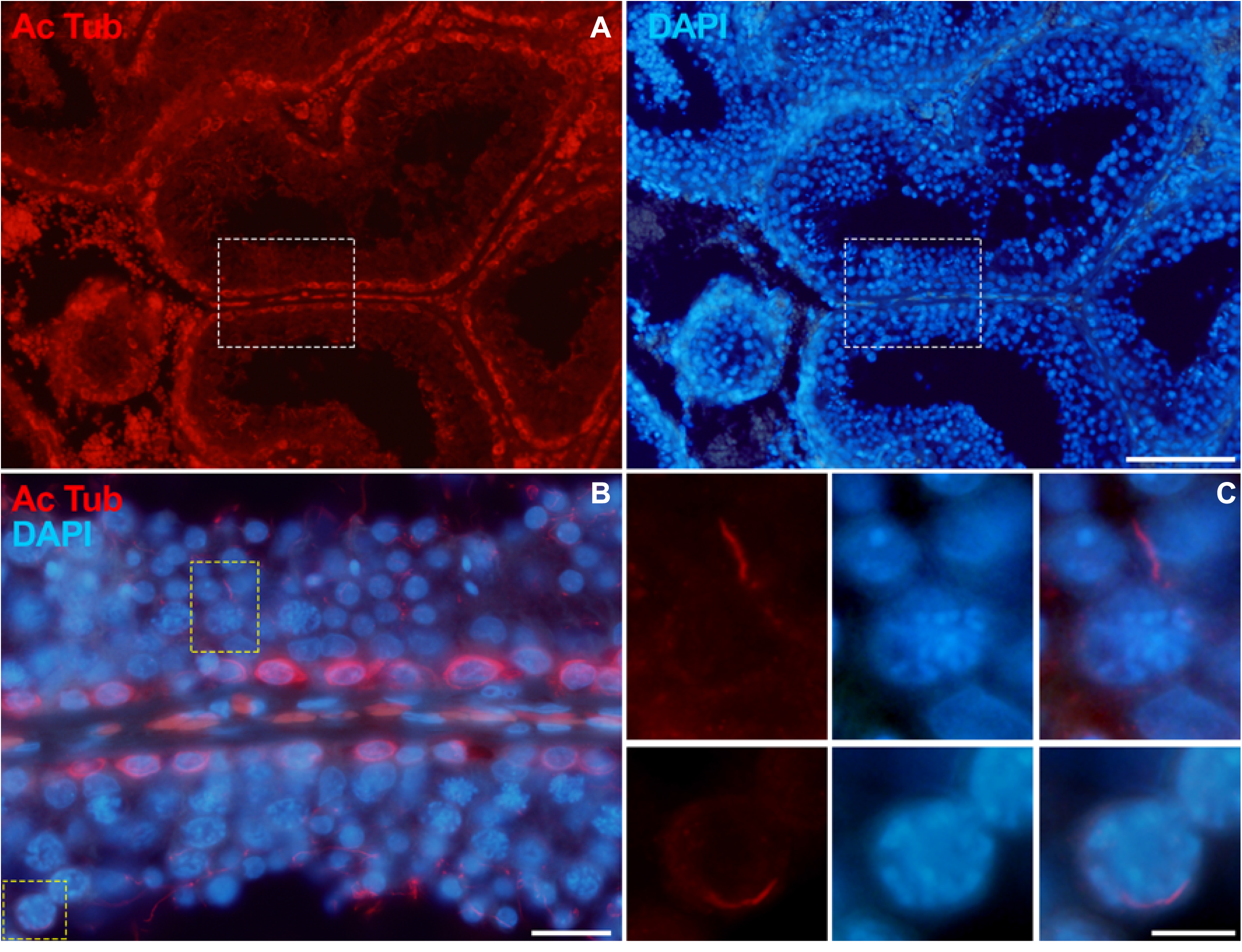
Human testis contains ciliated spermatocytes. **A-B.** Immunolabelling of acetylated Tubulin (AcTub) (red) in adult human testis cryosections. Chromatin is stained with DAPI (blue). **C.** Selected enhanced images correspond to two ciliated spermatocytes at the bases of the seminiferous tubule. Scale bars represent 50 µm in A, 20 µm in B and 10 µm in C.

**Supplementary Figure 5:**
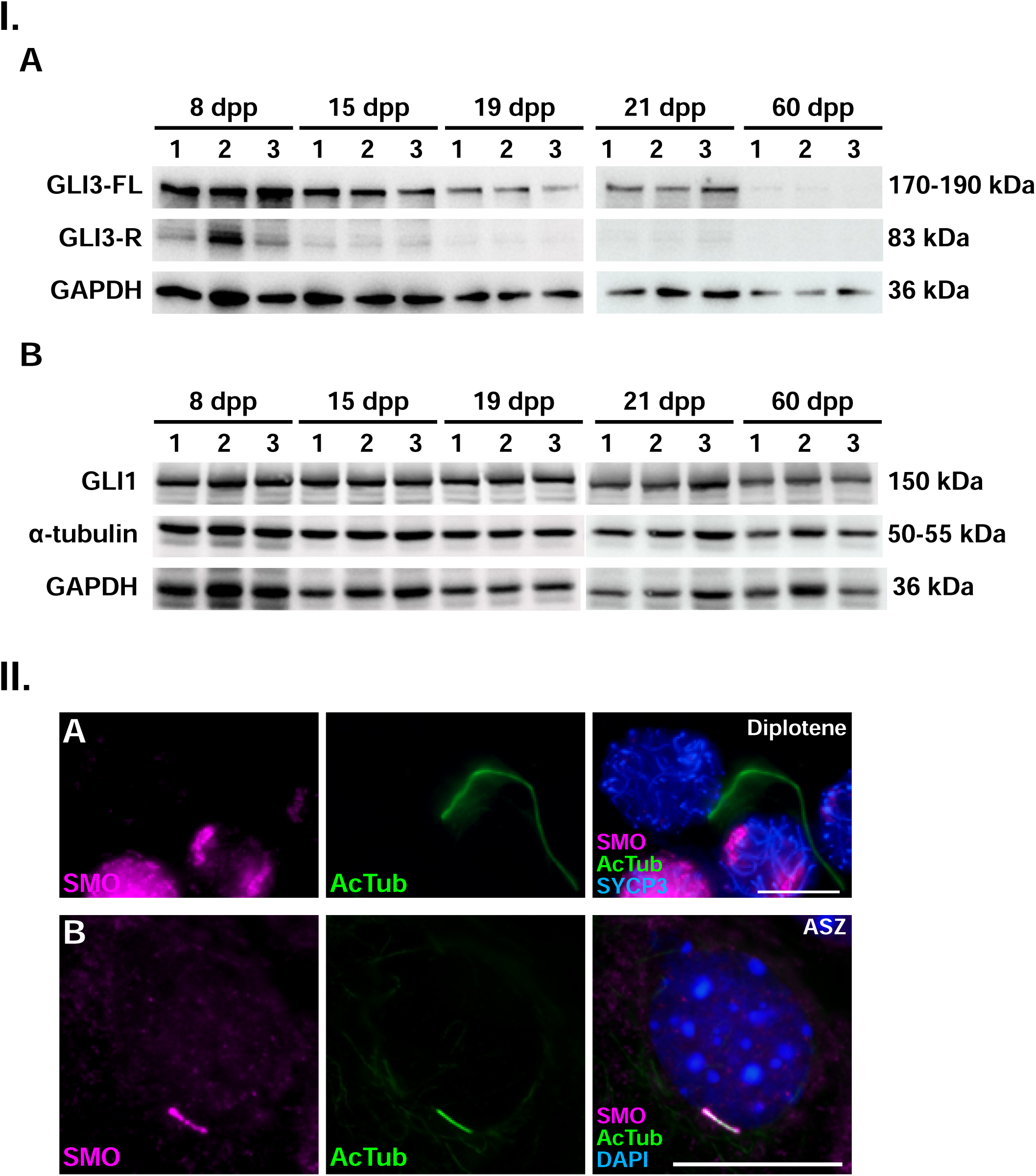
GLI1 and GLI3 readouts of Hedgehog signaling are not visibly affected in prepuberal mouse testes. **I.A.** Testis extracts from mice of various postnatal ages were resolved by SDS-PAGE and analyzed by Western blot using antibodies against GLI1, α-tubulin, and GAPDH. **B.** Testis extracts from the same developmental stages were analyzed using antibodies against GLI3 and GAPDH. The expected molecular weights of the target proteins are: GLI3 full-length (GLI3FL), 170–190 kDa; GLI3 repressor form (GLI3R), 83 kDa; GAPDH, 36 kDa.Mouse ages are indicated as postnatal days (8, 15, 19 and 21 dpp) or as adult (60dpp). Three biological replicates (n = 3) were analyzed for each age group. **II. A.** Double Immunolabelling of Smo (magenta) and acetylated Tubulin (AcTub) (green) in prepuberal 21 dpp spermatocytes. **II. B.** Double Immunolabelling of Smo (magenta) and acetylated Tubulin (AcTub) (green) in basal cell carcinoma cells. All images are z-projection of 30-60 frames of stack images of each cell. Chromatin is stained with DAPI (blue) in A, B. Scale bar represents 10 µm.

**Supplementary Figure 6:**
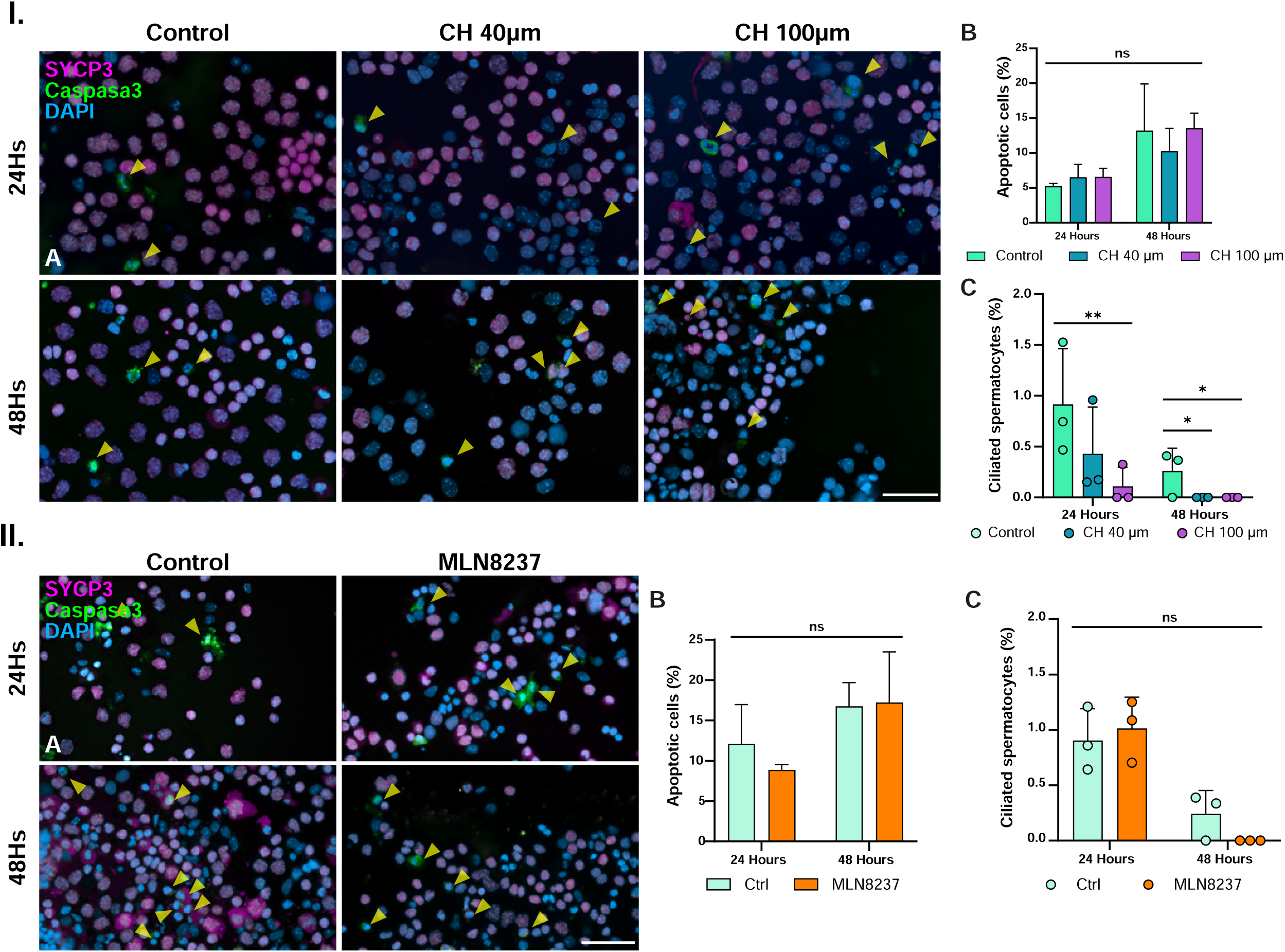
CH and MLN8237 optimization protocol. **I. Optimization of the chloral hydrate (CH) protocol. A.** Double immunolabelling of synaptonemal complex protein 3 (SYCP3) (magenta) and cleaved caspase 3 (green) in 40x magnification fields of cultured squashed spermatocyes from 20 dpp mice. Chromatin is stained with DAPI (blue). Experiments were conducted in 24-hours (24Hs) and 48-hours (48Hs) in control, 40 µM and 100 µM CH. Scale bar represents 10 µm. **B.** Graphical representation of the percentage of apoptotic spermatocytes for each condition of spermatocyte culture. Data represents no statistical significance (One-way ANOVA, p < 0.0001) (a minimum of 5500 spermatocytes per condition, three biological replicates). **C.** Graphical representation of the percentage of prophase I ciliated spermatocytes for each condition of spermatocyte culture. Data represents mean ± SD, **** p < 0.0001, Χ2 test (a minimum of 1500 spermatocytes per condition, three biological replicates. **II. Optimization of the MLN8237 protocol. A.** Double immunolabelling of synaptonemal complex protein 3 (SYCP3) (magenta) and cleaved caspase 3 (green) in 40x magnification fields of cultured squashed spermatocyes from 20 dpp mice. Chromatin is stained with DAPI (blue). Experiments were conducted in 24-hours (24Hs) and 48-hours (48Hs) in control and 10 µM MLN8237. Scale bar represents 10 µm. **B.** Graphical representation of the percentage of apoptotic spermatocytes for each condition of spermatocyte culture. Data represents no statistical significance (One-way ANOVA, p < 0.0001) (a minimum of 5500 spermatocytes per condition, three biological replicates). **C.** Graphical representation of the percentage of prophase I ciliated spermatocytes for each condition of spermatocyte culture. Data represents mean ± SD, **** p < 0.0001, Χ2 test (a minimum of 1500 spermatocytes per condition, three biological replicates.

**Supplementary Figure 7:**
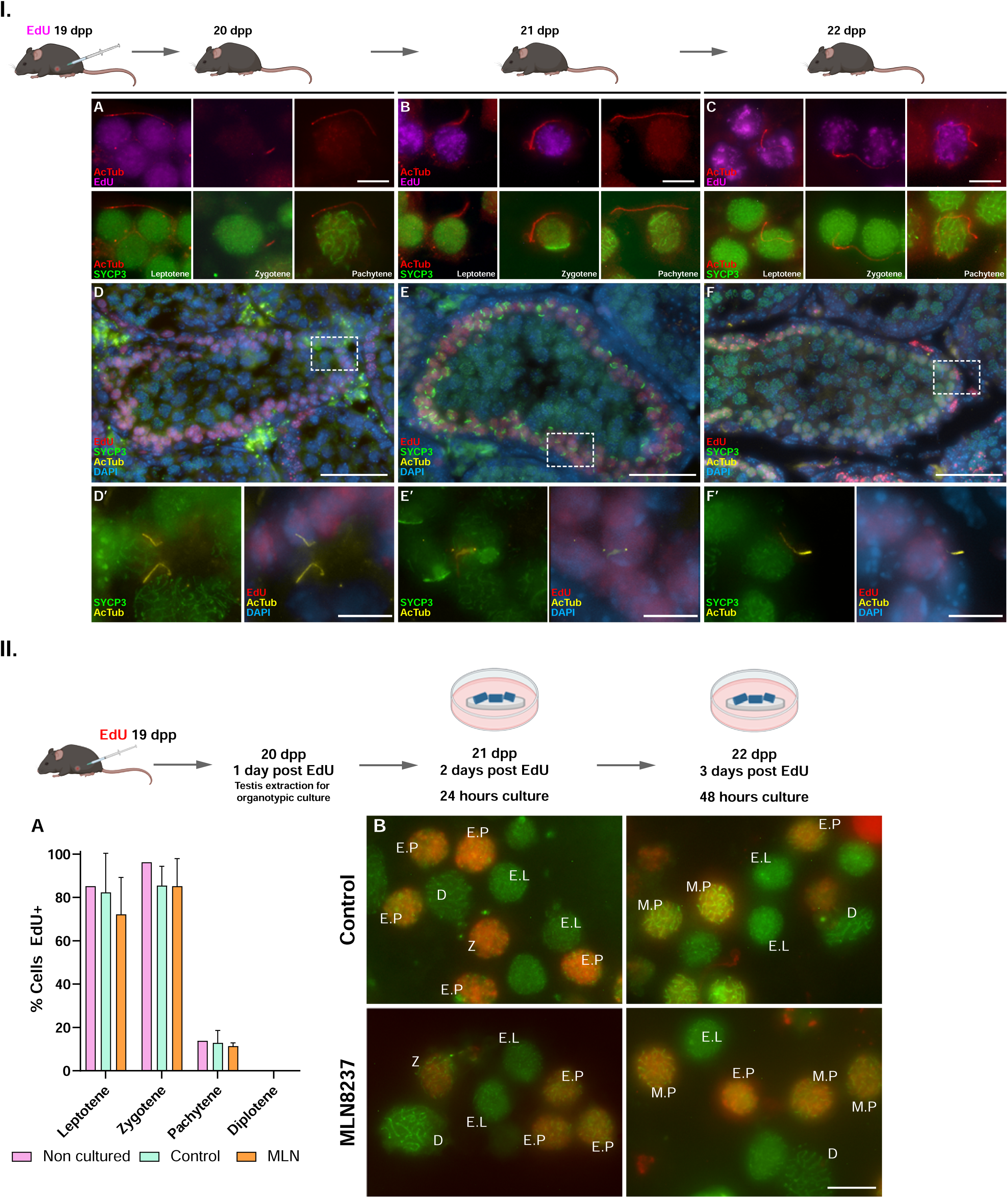
Tracking first meiosis wave with Edu pulse injection during prepubertal meiosis. **I.** Schematic representation of the protocol of EdU pulse. Injection of EdU is administered administered at 19 days post partum (dpp), followed by sacrifice of the animal an extraction of the testis after 24, 48 or 72 hours. **A-C.** Double immunolabelling of synaptonemal complex acetylated tubulin (AcTub) (red) and EdU (magenta) or synaptonemal complex protein 3 (SYCP3) (green) in squashed spermatocytes at prophase I stages (leptotene, zygotene and pachytene). Ciliated spermatocytes are presented for each stayed being Edu or non-Edu labelled, depending of the time of testis extraction. Scale bars represent 10 µm. **D-F.** Cryosection of the testis for timeframes 24, 48 or 72 hours after Edu injection. Triple immunolabeling of EdU labelling (red), synaptonemal complex protein 3 (SYCP3) (green) and acetylated tubulin (AcTub) (yellow). Chromatin is stained with DAPI (blue). Scale bars represent 10 µm. D’-F’. A magnified image of ciliated spermatocytes located at the base of the seminiferous tubules are presented. Scale bars represent 10 µm **II.** Schematic representation of the protocol of EdU pulse and posterior organotypic culture of seminiferous tubules. Injection of EdU is administered at 19 days post partum (dpp), followed by sacrifice of the animal an extraction of the testis after 24 hours, and posterior culture of fragments of seminiferous tubules for additional 24 or 48 hours. **A.** Graphical representation of the percentage of prophase I labelled with EdU after injection at 19 dpp and posterior incubation of spermatocytes during 24Hs in control and 10 µM MLN8237 samples. Data represents mean ± SD, Χ2 test (a minimum of 300 spermatocytes were quantified per condition, three biological replicates). **B.** Double immunolabeling of EdU labelling (red) and SYCP3 (green) in 40x fields of prophase I spermatocytes for control and MLN8237 condition. Each is labelled with its corresponding stage (E.L: early leptotene, Z: zygotene, E.P: early pachytene; M.P.: mid pachytene; D. Diplotene). Notice that given the absence of blood flow in the culture, there is no new incorporation of EdU in early leptotenes that are initiating meiosis. Scale bars represent 10 µm.

**Supplementary Figure 8:**
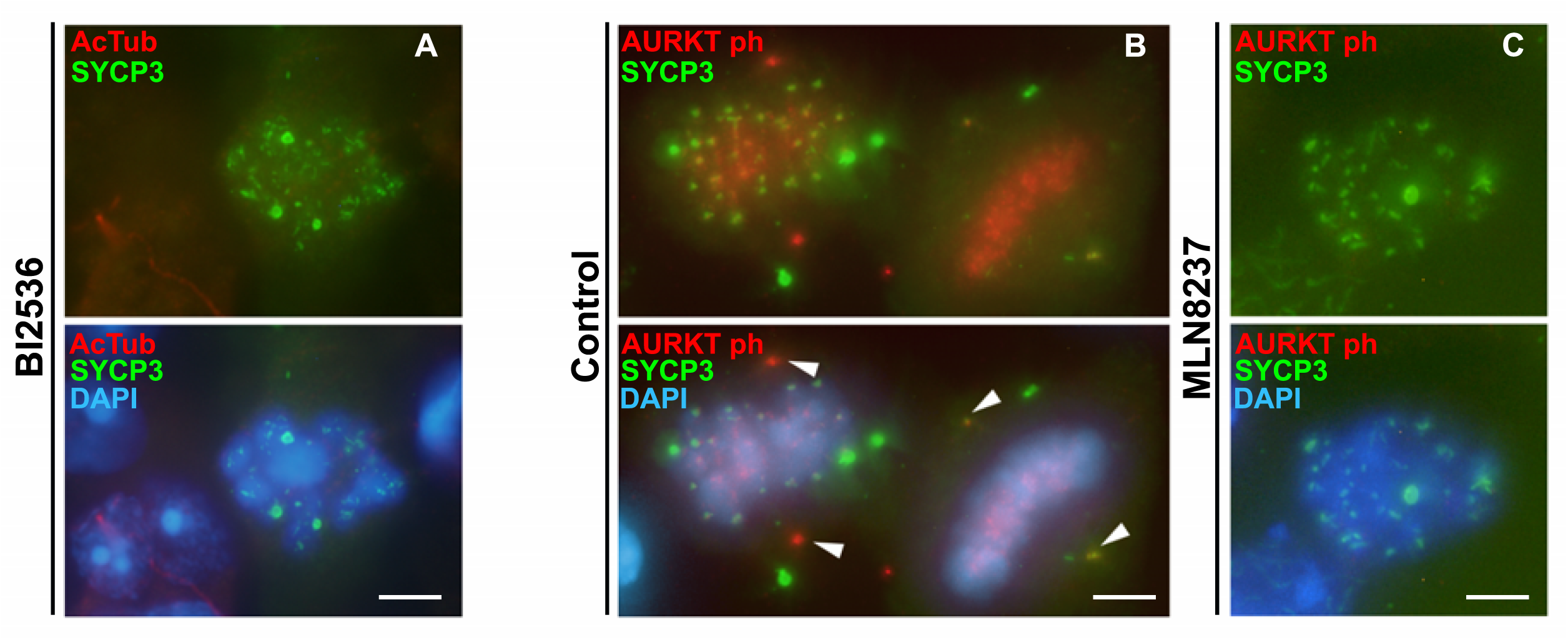
PLK1 and AURKA inhibition in cultured spermatocytes. **A.** PLK1 inhibition with BI2536 following previous reports 73. Double immunolabelling of acetylated tubulin (AcTub) (red) and synaptonemal complex protein 3 (SYCP3) (green) in a monopolar metaphase I. **B.** Double immunolabelling of phosphorylated AURK (AURKTph) 77 in control bipolar metaphase I and metaphase II. **C.** AURKA inhibition with MLN8237 following previous reports 73. Double immunolabelling of phosphorylated AURKTph in a monopolar metaphase I. Scale bars represent 10 µm.

